# CiliaQ – a simple, open-source software for automated quantification of ciliary morphology and fluorescence in 2D, 3D, and 4D images

**DOI:** 10.1101/2020.09.28.317065

**Authors:** Jan Niklas Hansen, Sebastian Rassmann, Birthe Stüven, Nathalie Jurisch-Yaksi, Dagmar Wachten

## Abstract

Cilia are hair-like membrane protrusions that emanate from the surface of most vertebrate cells and are classified into motile and primary cilia. Motile cilia move fluid flow or propel cells, while also fulfilling sensory functions. Primary cilia are immotile and act as a cellular antenna, translating environmental cues into cellular responses. Ciliary dysfunction leads to severe diseases, commonly termed ciliopathies. The molecular details underlying ciliopathies and ciliary function are, however, not well understood. Since cilia are small subcellular compartments, imaging-based approaches have been used to study them. However, tools to comprehensively analyze images are lacking. Automatic analysis approaches require commercial software and are limited to 2D analysis and only a few parameters. The widely used manual analysis approaches are time consuming, user-biased, and difficult to compare. Here, we present CiliaQ, a package of open-source, freely-available, and easy-to-use ImageJ plugins. CiliaQ allows high throughput analysis of 2D and 3D, static or time-lapse images from fluorescence microscopy of cilia in cell culture or tissues, and outputs a comprehensive list of parameters for ciliary morphology, length, bending, orientation, and fluorescence intensity, making it broadly applicable. We envision CiliaQ as a resource and platform for reproducible and comprehensive analysis of ciliary function in health and disease.

## 1 Introduction

Cilia are membrane protrusions that extend from the surface of almost all vertebrate cells. Cilia can either be motile or non-motile. Motile cilia generate a fluid-flow or propel cells forward, but also fulfil sensory function. Motile ciliated cells can bear a single, motile cilium, e.g. cells in the left-right organizer [1] or in the central canal of the spinal cord [2], or can bear bundles of cilia [3], e.g. cells with ependymal cilia in the brain [4, 5] or multi-ciliated cells in the respiratory tract [6]. Motile cilia can also be highly specialized, i.e. the sperm flagellum [7-10]. Non-motile, primary cilia function as cellular antennae that translate sensory information into a cellular response [11, 12]. Ciliary dysfunction leads to severe diseases commonly referred to as ciliopathies [13-15]. Patients display symptoms ranging from obesity, polycystic kidneys, or blindness to neurodevelopmental defects. Understanding the molecular details underlying the development of ciliopathies is crucial to develop treatment strategies.

To this end, it is important to characterize ciliary function and dysfunction with great level of detail and in a non-biased, automated fashion. Because most cilia are tiny compared to the rest of the cell, the analysis of ciliary function is image-based and involves parameters based on cilia morphology or protein localization, the latter determined using indirect fluorescent read-outs.

For the study of flagella, e.g., of sperm or algae, many analysis approaches have been established (e.g., [16-23]). Our group has released an open-source software for automated, comprehensive characterization of motile cilia and flagella based on 2D light and fluorescence microscopy imaging [24]. Similar software solutions have been released by other groups [25, 26]. However, such analysis approaches are not applicable for cilia in tissues and tissue culture. First, cilia on cells in a tissue are much smaller than flagella. Second, these cells cannot be analyzed solitarily, like sperm or algae. In turn, it is challenging to image an individual cilium in an intact tissue using light or 2D fluorescence microscopy, as a view on the cilium is precluded by other larger structures like the cell soma or by other cell layers. Third, flagellated, solitary cells, in contrast to ciliated tissue cells, can be tethered in an orientation that allows a simple analysis of the cilium. Albeit a similar approach has been established for live-cell imaging of tissue cells with fluorescently labeled cilia [27], its application is not trivial and not suited to study cilia in an intact tissue. Fourth, analyzing large numbers of cilia in the intact tissue is key to further understand the role of cilia, as cilia can act as a population and may require a specific orientation for proper functioning [28-30]. This is impractical if not impossible to establish with the available tools for flagella. In conclusion, analyzing large numbers of cilia in a tissue necessitates 3D imaging, fluorescent labels, and alternate image analysis approaches.

For analysis of cilia in tissues or tissue culture, most studies in the cilia field employ custom, mostly manual analysis of ciliary parameters, which makes it difficult to reproduce data in different labs and compare data sets from different studies. A previous study presented a software, revealing ciliary frequency (fraction of ciliated cells) and length with high throughput [31]: the software ACDC (<u>a</u>utomated <u>c</u>ilia <u>d</u>etection in <u>c</u>ells) detects and measures nuclei and primary cilia in 2D microscopy images. However, the software does not allow to analyze 3D images, relies on commercial software, and generates only a small number of parameters, which does not allow to extensively characterize cilia under physiological and pathological conditions. This is also true for other approaches to automatically analyze cilia, which relied on commercial software and were limited to either quantifying ciliary length [32, 33] or to determining ciliary distribution and orientation in a specific tissue [28].

To overcome all these limitations and foster reproducibility and comparability, we have developed CiliaQ, a software that allows automatized 3D reconstruction and comprehensive quantification of fluorescently-labeled cilia in 2D, 3D, or 4D images. CiliaQ quantifies a comprehensive list of parameters, allows batch-processing, time-lapse analysis, and correction of segmentations by the user, allowing high reproducibility, high throughput, and broad applicability. We exemplify CiliaQ analysis using microscopy images from immune-stained cultured cells and whole zebrafish embryos. This allowed us to precisely quantify Smoothened localization during activation of the Sonic Hedgehog pathway, to delineate the differences of cilia labeling with an acetylated-Tubulin antibody, Arl13B antibody, or Arl13B-GFP expression, to demonstrate potential biases of 2D vs. 3D analysis, and to unravel that cilia in the zebrafish embryo are specifically oriented during neuronal delamination. Thereby, CiliaQ presents an unprecedented resource for analysis of cilia in the tissue.

## 2 Results

### 2.1 The CiliaQ analysis workflow

CiliaQ constitutes a three-step workflow based on three ImageJ plugins (Fig. 1A): (1) CiliaQ Preparator, which prepares the image for segmentation and segments the image into cilia and background, (2) CiliaQ Editor, which allows manual corrections of the segmentation, and (3) CiliaQ, which fully-automatically reconstructs the cilia, quantifies them, and visualizes the results. The workflow requires 3D confocal image stacks or 2D fluorescence images as input, which need to contain one channel with a cilia marker (cilia mask) to allow cilia reconstruction. Additional channels may label other proteins of interest, whose ciliary localization can be studied using CiliaQ. Of note CiliaQ is not limited to static 3D images. Although not presented in this paper, CiliaQ can also be employed to analyze time-lapse 3D images. A CiliaQ analysis using data from a spinning-disk confocal microscope has recently been published [34].

**Figure 1:**
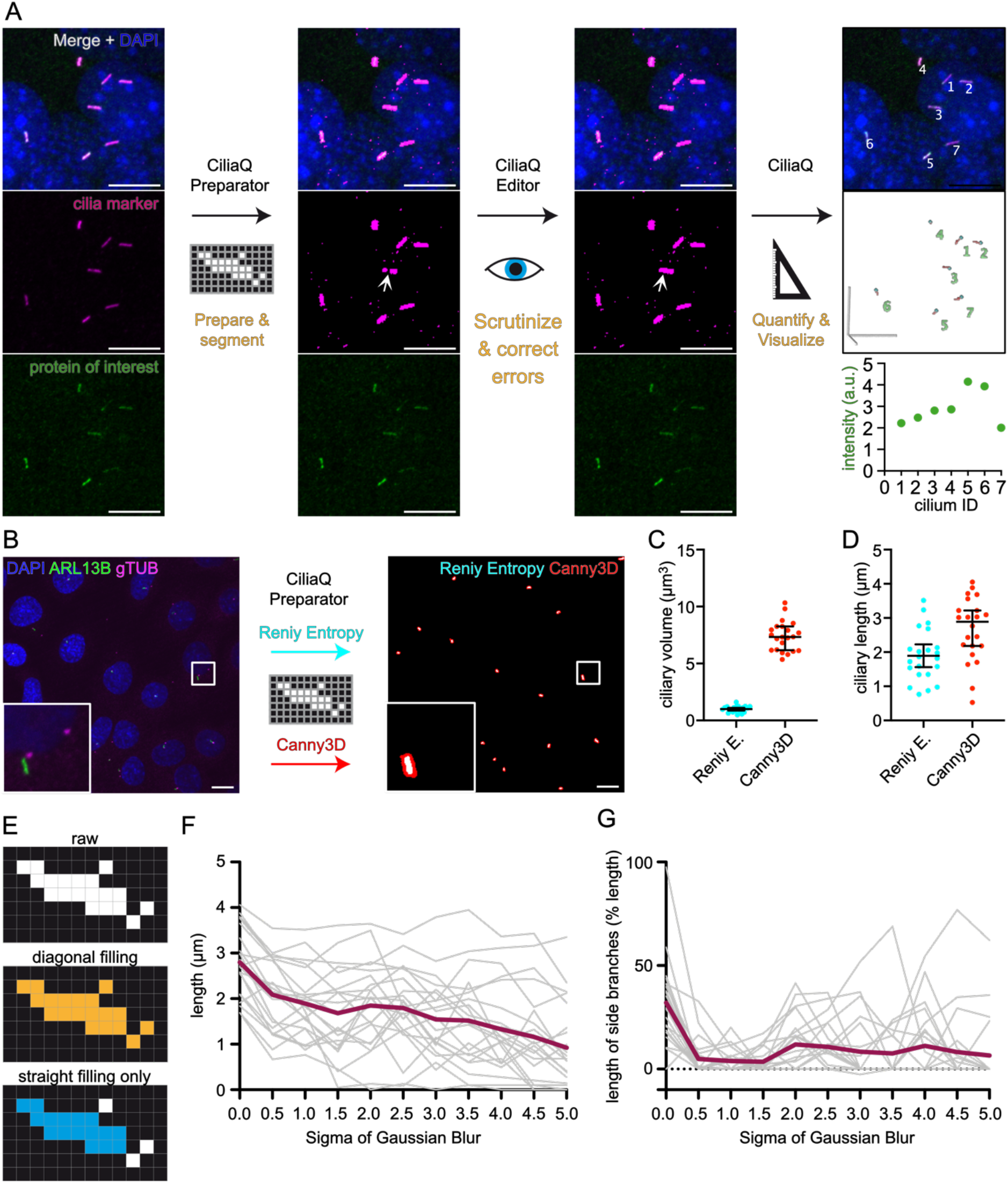
CiliaQ workflow. **(A)** CiliaQ constitutes a three-step workflow based on three ImageJ plugins: CiliaQ Preparator, which automatically segments the image into ciliary and background voxels, CiliaQ Editor, which allows manual correction of segmentation errors, and CiliaQ, which automatically reconstructs and quantifies all cilia in the image and outputs and visualizes the results. **(B-D)** Comparing segmentation with a single intensity threshold, determined by a histogram-based threshold method (Reniy Entropy), or with the Canny3D method in CiliaQ Preparator for processing an exemplary confocal stack image acquired from fixed, serum-deprived wild-type IMCD-3 cells, stained with DAPI to label nuclei (blue), an ARL13B antibody to label cilia (green), and a gamma-Tubulin antibody (gTUB) to label centrosomes (magenta). **(B)** Maximum projection of the original stack image (left) and the ARL13B channel after segmentation with CiliaQ Preparator using the Reniy-Entropy threshold (cyan) or the Canny3D method (red) (right). Magnified view on lower left. **(C)** Ciliary volume and **(D)** ciliary length determined by CiliaQ analysis of the segmented image stack shown in B using either the Reniy-Entropy-threshold- or the Canny3D-based segmentation for ciliary reconstruction. Each data point represents an individual cilium. All scale bars: 10 µm. **(E)** Detecting objects in a pixelated binary image (top) with a Flood-Fill-Algorithm filling in horizontal, vertical, and diagonal direction (orange, center) or filling in horizontal and vertical direction only (blue, bottom). **(F-G)** Dependency of length quantification on the Gaussian blur applied (sigma in pixels) prior to skeletonization. **(F)** Length of the cilia presented in B (Reniy-Entropy threshold) determined by CiliaQ analysis with different Gaussian Blur sigma settings. **(G)** Total length of side-branches appearing in the skeleton generated for measuring the ciliary length, presented as % length of the main branch.

#### 2.1.1 CiliaQ Preparator

After specifying the processing settings and the channel for the cilia mask, CiliaQ Preparator fully automatically segments the image into cilia and background. For pre-processing, CiliaQ Preparator offers background subtraction, which is particularly helpful when analyzing images acquired with unequal illumination or to remove signal from background structures larger than cilia.

For segmentation, CiliaQ Preparator offers standard intensity threshold algorithms using a hysteresis threshold, and a custom developed method that we coined “Canny3D”, as it is based on a 3D implementation of Canny edge detection [35]. The user can choose whether threshold algorithms use the stack’s intensity histogram or the histogram of a maximum intensity projection of the stack for calculation. The latter advantageously increases the relationship of fore-to background in 3D images of cilia that commonly contain little foreground compared to background pixels. The Canny3D method employs four consecutive steps: (1) the image is smoothed with a 2D Gaussian kernel applied to each slice, (2) edges are detected with a 3D Sobel kernel, (3) a 3D Hysteresis threshold is applied, and (4) holes encapsulated in all three dimensions are filled. For steps (2) to (4), we make use of functions from the ‘3D ImageJ Suite’ [36], an open-source software extension for ImageJ.

Each method for segmentation is applicable and features advantages and disadvantages. For example: Canny3D detects the edges of the cilium and not the cilium itself, whereby it generates bigger segmentations of cilia and thereby, reduces detection gaps in incompletely labeled cilia (Fig. 1B). Accordingly, parameters quantified by CiliaQ, e.g. ciliary volume (Fig. 1C) or ciliary length (Fig. 1D) differ between data analyzed with the Canny3D method and threshold-based segmentation methods. Generally, the segmentation method needs to be adapted to the experimental paradigm and thus, for each experimental paradigm, the settings need to be optimized before analysis. However, for all images of a connected data set, the same settings need to be used. Low-performing segmentation paradigms or non-optimized settings may require extensive manual corrections in the second step of the CiliaQ analysis pipeline.

#### 2.1.2 CiliaQ Editor

CiliaQ Editor is a manual tool that allows to correct the segmentation in the output image of CiliaQ Preparator while documenting every correction step.

#### 2.1.3 CiliaQ

CiliaQ reconstructs the ciliary objects based on the image prepared by CiliaQ Preparator and, eventually, CiliaQ Editor. Initially, the channel that contains the segmented cilia (reconstruction channel) needs to be specified. Optionally, additional channels can be specified, in which the fluorescence intensity in the cilium has to be measured (channel labeling a protein of interest) or which contain a staining of the ciliary base (basal stain). The latter allows CiliaQ to orient intensity profiles along the cilium from base to tip.

To reconstruct cilia, CiliaQ detects objects in the image using a 3D Flood-Filling-Algorithm on the segmented channel, for which two variants are offered. They differ by Flood-Filling in all directions (horizontal, vertical, and diagonal, Variant 1) or in the horizontal and vertical direction only (Variant 2) (Fig. 1E). While variant 1 better retrieves incompletely labeled cilia, this procedure may include more noise into ciliary objects, especially in images with low signal-to-noise ratio. Furthermore, variant 1 may more frequently connect adjacent cilia to one ciliary object in densely ciliated images.

After reconstruction, cilia objects are filtered by a user-defined size threshold. Cilia below threshold are considered as noise and excluded. In addition, CiliaQ offers means to exclude cilia that are incompletely detected because they are localized at image borders. To fully capture cilia, it is recommended to exclude cilia touching x, y, or, z borders in a 3D analysis and exclude cilia touching x or y borders in a 2D analysis.

To determine length, intensity profiles, tangent vectors, and curvature profiles of cilia, CiliaQ uses the ImageJ plugins Skeletonize3D_ and AnalyzeSkeleton_ [37]. CiliaQ skeletonizes each cilium and extracts the centerline from the largest shortest path of the skeleton. The largest shortest path is determined as follows: for all combinations of two end-points of the skeleton, the shortest connecting path on the skeleton is determined. A combination of two-endpoints with the longest connecting path is selected and the path in-between represents the largest shortest path in the skeleton. To improve the skeletonization of the ciliary object similarly to what has been described [33], the following image processing is applied: a 3-fold upscaled image of the ciliary object is generated and blurred with a Gaussian blur. The sigma of the blur is user-defined and may be adjusted depending on the noise level in the image. Increasing the sigma will remove more noise prior to skeletonization, but also generally shortens the measured length (Fig. 1F). While a Gaussian blur with a low sigma reduces the appearance of side branches, at higher sigma values, the appearance of side branches in the skeletonization increases (Fig. 1G), which in turn may cause imprecise measurements of the ciliary length. For example, for the presented setting (Fig. 1B), Gaussian blur sigma settings of 0.5 to 1.5 pixel revealed the minimal generation of side branches for all cilia in the image (Fig. 1G).

CiliaQ quantifies morphology (Fig. 2A-E) and intensity (Fig. 2F-K) parameters. The intensity-based parameters are determined for the user-specified channels labeling a protein of interest. Intensity-based parameters not requiring the intensity profile (average, minimum, maximum, standard deviation of fluorescence intensity (Fig. 2F) and colocalized volume (Fig. 2J)) are also reported for the reconstruction channel. A complete list of all parameters is shown in Table 1.

**Table 1:** CiliaQ output parameters. Some parameters are output in multiple ways with different units. For 2D analysis, the numeric values of volume parameters correspond to the area of the cilium in the image. The parameters Shape complexity index and Sphere radius are not applicable for 2D analysis. The parameters Arc length, Tangent vector, and Signed 3D Curvature are output for each individual point on the ciliary centerline. Abbreviations: a.u. (arbitrary units), voxel (a pixel in a 3D image).

| Parameter | Units | Description |
| --- | --- | --- |
| x-, y-, z-center | $\mu\text{m}$ | Center of the cilium |
| Volume | voxel,<br>$\mu\text{m}^3$ | The volume of the cilium as number of voxels or metric volume (calculated as the product of the number of voxels, the voxel width, the voxel height and the voxel depth). |
| # surface voxels | - | The number of voxels belonging to the cilium that are localized at the ciliary surface |
| Surface | $\mu\text{m}^2$ | The surface of the cilium. Determined as the sum of the individual voxel surfaces with each voxel's surface defined as the border surfaces of the voxel i, at which no neighbored ciliary voxel exists |
| Shape complexity index | - | Comparing the surface of the cilium to the surface of a perfect sphere containing the same volume of the cilium, this parameter gives a measure on how aspherical the cilium's shape is.<br>Determined as $\frac{\text{surface}}{4\pi \cdot ((3 \cdot \text{volume}) / (4\pi))^{2/3}}$. An index of 1 represents a perfectly spherical shape. The higher the index, the less spherical the shape |
| Sphere radius | $\mu\text{m}$ | The radius of a sphere containing the same volume as the cilium |
| A: Colocalized volume | $\mu\text{m}^3$,<br>% of<br>total<br>volume | The volume of the cilium that is non-zero in channel A (only applicable if the intensity channel A is background-removed / binarized) |
| B: Colocalized volume | $\mu\text{m}^3$,<br>% of<br>total<br>volume | The volume of the cilium that is non-zero in channel B (only applicable if the supplied channel B is background-removed / binarized) |
| A: Colocalized compared to BG volume | $\mu\text{m}^3$,<br>% of<br>total<br>volume | The ciliary enrichment of a stained or fluorescent protein (labeled in channel A) compared to the expression level in the soma (BG volume = Background volume). Determined as the ciliary volume with higher intensity than a threshold defined by the soma intensity. The threshold is calculated as follows:<br>Divide the entire image into 25 equally-sized cuboids |
|  |  | Collect from each cuboid the 10% of voxels that feature the highest intensities in the cuboid but that are not part of any cilium object.<br>Threshold = Average + 1.5-fold standard deviation of the intensities of the voxels collected from all cuboids |
| B: Colocalized compared to BG volume | $\mu\text{m}^3$,<br>% of<br>total<br>volume | The ciliary enrichment of a stained or fluorescent protein (labeled in channel B) compared to the expression level in the soma (BG volume = Background volume). Determined as the ciliary volume with higher intensity than a threshold defined by the soma intensity. The threshold is calculated as follows:<br>Divide the entire image into 25 equally-sized cuboids<br>Collect from each cuboid the 10% of voxels that feature the highest intensities in the cuboid but that are not part of any cilium object.<br>Threshold = Average + 1.5-fold standard deviation of the intensities of the voxels collected from all cuboids |
| minimum intensity (in reconstruction channel, channel A, or channel B) | a.u. | Minimum intensity of the voxels contained in the ciliary volume. |
| maximum intensity (in reconstruction channel, channel A, or channel B) | a.u. | Maximum intensity of the voxels contained in the ciliary volume. |
| average intensity (in reconstruction channel, channel A, or channel B) | a.u. | Average intensity of the voxels contained in the ciliary volume. |
| average intensity of the 10% of voxels with highest intensity (in reconstruction channel, channel A, or channel B) | a.u. | Average intensity of the 10% of all voxels contained in the ciliary volume with the highest intensities in the cilium. |
| SD of intensity (in reconstruction channel, channel A, or channel B) | a.u. | Standard deviation (SD) of all voxels' intensities in the ciliary volume. |
| # of found skeletons (quality parameter) | - | If above 1, indicative for incomplete or incorrect cilium reconstruction. In turn, all skeleton-based parameters might be incorrect. |
| # branches (quality parameter) | - | If above 1, the detected ciliary skeleton might contain side branches, which can be indicative for an incorrect ciliary detection. |
| tree length (quality parameter) | $\mu\text{m}$ | If largely different from the parameter "cilia length", can be indicative for an incorrect ciliary detection. |
| cilia length | μm | The arcus length of the cilium. Determined as the largest shortest path of the largest skeleton detected for the cilium. |
| orientation vector x, y, and z | μm | The orientation of the cilium in space. Determined as the vector from first (cilium base) to last (cilium tip) skeleton point. |
| cilia bending index | - | The bending / curvature of the cilium. Determined as the arcus length along the cilium divided by the Euclidian distance of first (cilium base) and last (cilium tip) skeleton point. |
| Intensity threshold A | a.u. | The threshold determined to calculate the parameters “Colocalized compare to BG volume” and “Colocalized on centerline compared to BG volume” in channel A.<br>The threshold is calculated as follows:<br>• Divide the entire image into 25 equally-sized cuboids<br>• Collect from each cuboid the 10% of voxels that feature the highest intensities in the cuboid but that are not part of any cilium object.<br>• Threshold = Average + 1.5-fold standard deviation of the intensities of the voxels collected from all cuboids |
| Intensity threshold B | a.u. | The threshold determined to calculate the parameters “Colocalized compare to BG volume” and “Colocalized on centerline compared to BG volume” in channel B.<br>The threshold is calculated as follows:<br>• Divide the entire image into 25 equally-sized cuboids<br>• Collect from each cuboid the 10% of voxels that feature the highest intensities in the cuboid but that are not part of any cilium object.<br>• Threshold = Average + 1.5-fold standard deviation of the intensities of the voxels collected from all cuboids |
| Integrated A intensity |  | The total intensity level in channel A along the ciliary centerline, determined as the sum of intensities along the largest shortest path of the detected skeleton. |
| Average A intensity on centerline |  | The average intensity level in channel A along the ciliary centerline, determined as the average of intensities along the largest shortest path of the detected skeleton. |
| Integrated B intensity |  | The total intensity level in channel B along the ciliary centerline, determined as the sum of intensities along the largest shortest path of the detected skeleton. |
| Average B intensity on centerline |  | The average intensity level in channel B along the ciliary centerline, determined as the average of intensities along the largest shortest path of the detected skeleton. |
| A: Colocalized on centerline compared to BG volume | $\mu\text{m}$,<br>% total<br>length | The ciliary enrichment of a stained or fluorescent protein (labeled in channel A) compared to the expression level in the soma (BG volume = Background volume). Determined as the length/percentage of the ciliary centerline (the largest shortest path of the detected skeleton) featuring intensities above a threshold defined by the soma intensity.<br>The threshold is calculated as follows:<br>• Divide the entire image into 25 equally-sized cuboids<br>• Collect from each cuboid the 10% of voxels that feature the highest intensities in the cuboid but that are not part of any cilium object.<br>Threshold = Average + 1.5-fold standard deviation of the intensities of the voxels collected from all cuboids |
| B: Colocalized on centerline compared to BG volume |  | The ciliary enrichment of a stained or fluorescent protein (labeled in channel B) compared to the expression level in the soma (BG volume = Background volume). Determined as the length/percentage of the ciliary centerline (the largest shortest path of the detected skeleton) featuring intensities above a threshold defined by the soma intensity.<br>The threshold is calculated as follows:<br>• Divide the entire image into 25 equally-sized cuboids<br>• Collect from each cuboid the 10% of voxels that feature the highest intensities in the cuboid but that are not part of any cilium object.<br>• Threshold = Average + 1.5-fold standard deviation of the intensities of the voxels collected from all cuboids |
| Profile A | a.u. | The intensities in channel A along the ciliary centerline (the largest shortest path of the detected skeleton). For each point of the largest shortest path, the intensity is interpolated from the four neighboring pixels in the closest stack image. For output, the profile is scaled in steps of the voxel width, for which adjacent points are averaged. |
| Profile B | a.u. | The intensities in channel B along the ciliary centerline (the largest shortest path of the detected skeleton). For each point of the largest shortest path, the intensity is interpolated from the four neighboring pixels in the closest stack image. For output, the profile is scaled in steps of a voxel width, for which adjacent points are averaged. |
| Arc length | $\mu\text{m}$ | The arc length position of each point on the ciliary centerline (the largest shortest path of the detected skeleton). Defined by the distance of a given point to the first ciliary point on the centerline. |
| Tangent vector x, y, and z | $\mu\text{m}$ | The tangent vectors along the ciliary centerline (the largest shortest path of the detected skeleton). For each point of the largest shortest path, a tangent vector is determined as the vector from the point at a specific arc length distance upstream on the cilium to the point at a specific arc length distance downstream on the cilium (the specific distance is set by the user). The tangent vectors are normalized to a length of unity. |
| Signed 3D Curvature | $\mu\text{m}^{-1}$ | The curvature along the ciliary centerline (the largest shortest path of the detected skeleton). At each point on the cilium the curvature is determined based on the equation for the geometric curvature.<br>More specifically, the curvature is determined from the normalized tangent vectors of an upstream point P1 and downstream point P2 (tangent vectors T1 and T2, respectively) as: $(T2 - T1) / 2 / (\text{arc length distance P1 to P2 on the cilium})$. Finally, the curvature is signed by the sign of the cross product of T1 and T2. |

**Figure 2:**
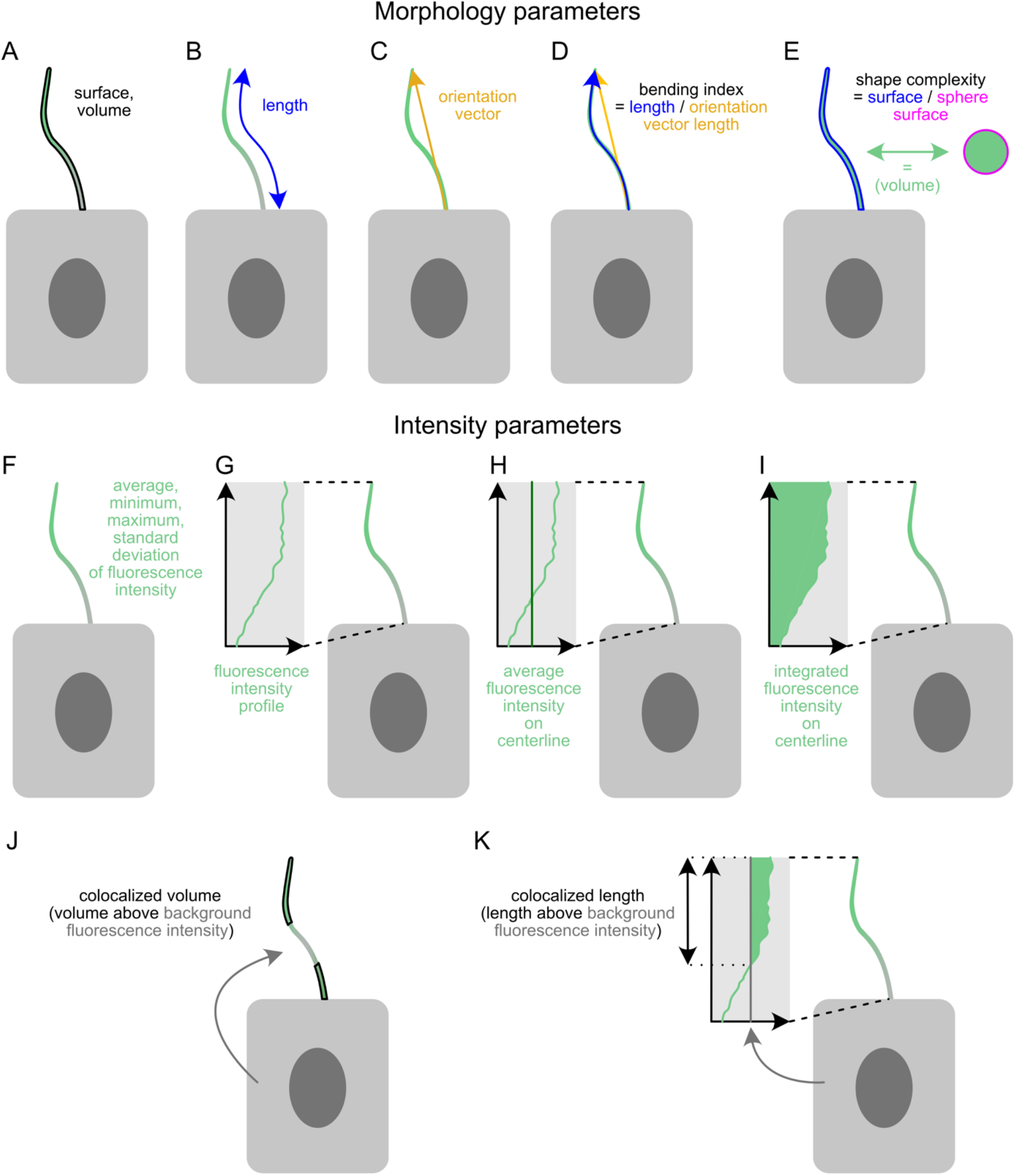
CiliaQ output parameters. For each reconstructed cilium, CiliaQ determines **(A-E)** structural parameters and **(F-K)** intensity parameters for up to two user-defined intensity channels. **(A)** Ciliary surface and ciliary volume. **(B)** Ciliary length. **(C)** Ciliary orientation vector, the vector pointing from the base to the tip of the cilium. **(D)** Ciliary bending index, determined as the ratio of ciliary length and the length of the orientation vector. **(E)** Ciliary shape complexity index, determined as the ratio of the ciliary surface and the surface of a sphere containing the ciliary volume. **(F)** Average, minimum, maximum, and standard deviation of the fluorescence intensity values of ciliary voxels. **(G)** Fluorescence intensity profile along the ciliary centerline. **(H)** Average fluorescence intensity along the ciliary centerline. **(I)** Integrated fluorescence intensity on the ciliary centerline. **(J)** Colocalized ciliary volume compared to background, determined as the ciliary volume comprising voxels with fluorescence intensities above an intensity threshold that is calculated based on the general background intensity level in the image. **(K)** Colocalized ciliary length, determined as the ciliary length in the intensity profile with intensities above an intensity threshold that is calculated as described for I.

Finally, CiliaQ also outputs 3D visualizations of the reconstructed cilia and the “largest shortest paths” in the ciliary skeleton, visualizing the detected ciliary length and intensity profile position and direction (see for example Fig. 1A).

#### 2.1.4 Subsequent data analysis

Data sets of cilia commonly contain many images. CiliaQ outputs an individual results file for each image, providing the parameters for each cilium in the image. Thus, after CiliaQ analysis, the results tables need to be joined in an efficient way to avoid exhausting manual copy-paste-steps. To facilitate the subsequent analysis of the results, we provide an R template (see User Guide) that allows to automatically extract CiliaQ results from the file system and merge the results. The script allows to easily produce results tables for specific parameters, plot the results, and perform further bioinformatic analysis of the parameters. For example, the template shows how to identify the parameters that mostly define differences between analyzed conditions using Principle Component Analysis or how to unravel correlated parameters for the population of analyzed cilia.

### 2.2 Analyzing ciliary protein localization with CiliaQ

To demonstrate an exemplary CiliaQ analysis of ciliary protein localization, we generated a data-set according to a commonly studied paradigm in cilia research and analyzed the data set using CiliaQ. More specifically, we analyzed serum-deprived mouse embryonic fibroblasts after Smoothened-agonist (SAG) treatment. SAG treatment stimulates Sonic hedgehog (Shh) signaling, and, thereby the accumulation of Smoothened (SMO) in the cilium [38]. Cilia and SMO were visualized using antibody labeling (Fig. 3A, B). We recorded two confocal image stacks per condition (control and SAG treatment), and submitted these image stacks to the CiliaQ analysis pipeline (Fig. 3C). Along the analysis, we tracked the time of user-dependent and -independent analysis (Fig. 3).

**Figure 3:**
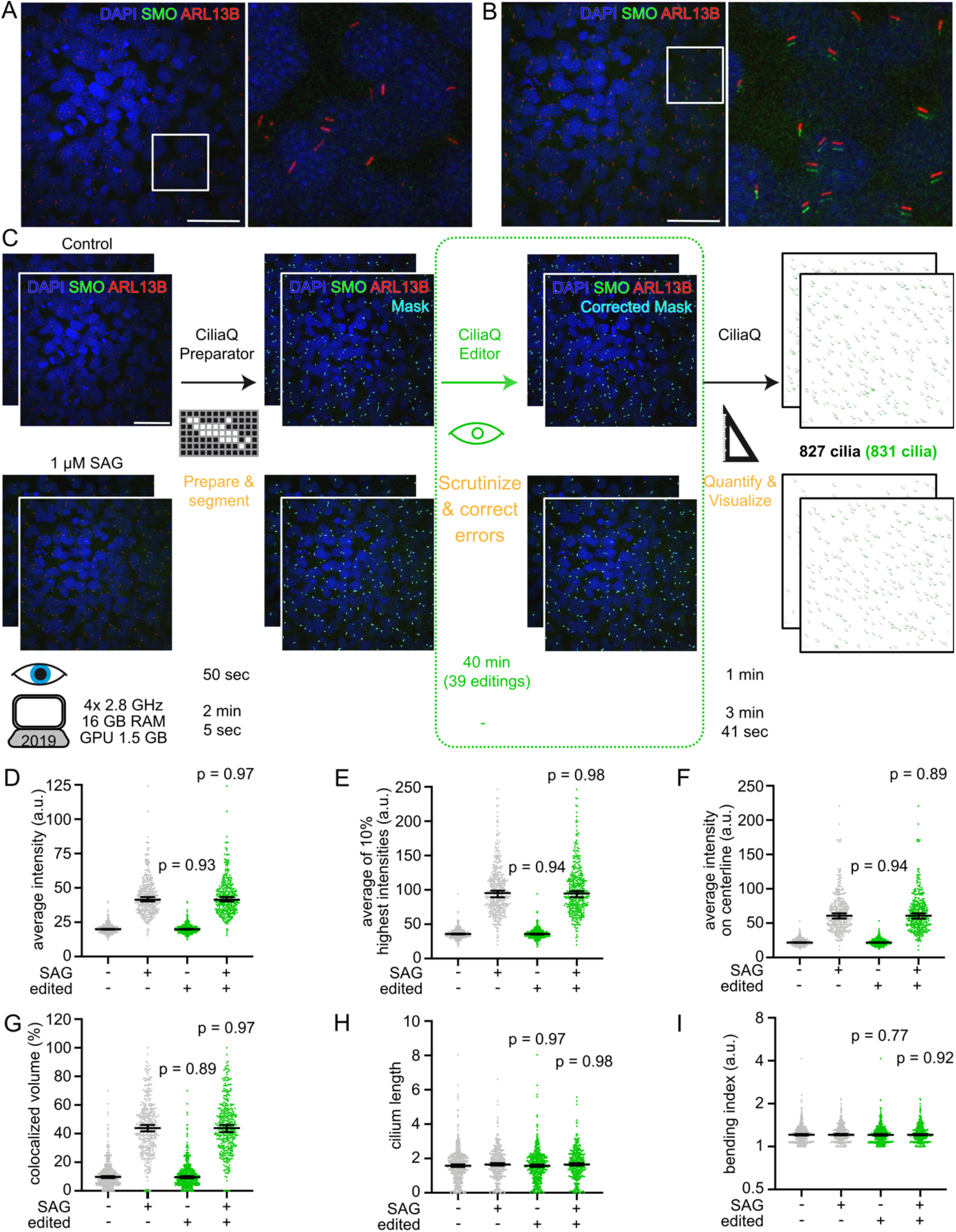
Exemplary CiliaQ analysis of ciliary localization of a protein of interest. **(A-B)** Exemplary confocal 3D stacks acquired from serum-starved mouse embryonic fibroblast cells that were either unstimulated (A) or stimulated with smoothened agonist (SAG, 1 µM, B) for 24 h before fixation and staining with an ARL13B antibody to label cilia, a Smoothened (SMO) antibody, and with DAPI to label nuclei. Scale bars: 50 µm. In all images, the green channel (SMO) was shifted by 7 px to the bottom for better visualizing SMO accumulation in cilia. **(C)** Analysis workflow for the data set presented in A and B, including two 3D stacks per condition. The four stacks were segmented using CiliaQ Preparator and next, either directly quantified with CiliaQ Editor (black steps applied only in workflow), revealing 827 cilia, or segmentation errors were corrected with CiliaQ Editor before quantification of the images with CiliaQ (black and green steps applied in workflow), revealing 831 cilia. Below the individual steps, the time required for the analysis is plotted: Eye: the time requiring direct user interaction; Computer symbols: the time for computer-controlled automated analysis without user-interaction on notebooks with the indicated specifications. Scale bar: 50 µm. **(D-I)** CiliaQ results of the analysis demonstrated in C. D to G describe the intensity of the SMO signal. H and I describe morphological parameters. Grey: CiliaQ results obtained from uncorrected segmented images. Green: CiliaQ results obtained from images corrected/edited with CiliaQ Editor. P-values indicate the test results of a Mann-Whitney test compared to the uncorrected results for the respective condition. Each data point represents an individual cilium. Bars indicate median ± 95% confidence interval of the median.

Segmenting the cilia from background with CiliaQ Preparator required only few minutes. Next, we either directly analyzed the segmented images with CiliaQ or scrutinized every individual cilium in the image prior to CiliaQ analysis using CiliaQ Editor. Thereby, we could assess the errors of a fast, fully automated cilia detection without manual corrections compared to a slower, semi-automated but high-fidelity cilia detection, involving manual correction with CiliaQ Editor. While the correction step requires user interaction (in this case about 10 min/image), CiliaQ – like CiliaQ Preparator – is fully automated and offers batch processing, allowing to run it as a background application. In total, CiliaQ analysis of the four images took about four minutes on a state-of-the-art notebook.

In the scrutinization step, we discovered and corrected 39 (5% in total) inaccuracies of the segmentation, where either multiple cilia were connected or where cilia were incompletely detected. This only reduced the total number of detected cilia by 0.5% (Uncorrected: 821 detected cilia; Corrected: 831 detected cilia).

CiliaQ quantified a significant increase in ciliary SMO signal in the different intensity parameters, such as the average intensity (Fig. 3D), average intensity of the 10% highest pixels in the cilium (Fig. 3E), the average intensity on the centerline (Fig. 3F), and the colocalized volume compared to the background (Fig. 3G). The different intensity parameters imply different detection biases. Generally, the average of the 10% highest intensity pixels in the cilium (Fig. 3E) and the average intensity on the centerline (Fig. 3F) are less sensitive to errors introduced during the segmentation process due to a low signal-to-noise-ratio. For the parameter “average of the 10% highest intensity pixels in the cilium”, only the brightest ciliary pixels are considered, excluding noise. However, this parameter depends on a uniform ciliary intensity distribution or only reflects the intensity level of the brightest spots in the cilium. In contrast, the average intensity on the centerline is independent of the intensity distribution along the cilium and does not include the intensities at the ciliary border, where the ciliary mask can be subject to noise, thereby, representing a more stable analysis parameter than the average ciliary intensity. The parameter “colocalized volume compared to background” normalizes the ciliary intensity to the intensity in the somas of all cells in the image and, thereby, gives a measure for the volumetric ciliary protein enrichment compared to the cell soma (Fig. 3G). For example, the median of ciliary SMO colocalization was about 10% at basal conditions and about 40% after SAG stimulation.

Notably, the general outcome of the experiment remained unchanged by leaving out the CiliaQ Editor (Fig. 3D-G). Not only intensity parameters, but also the results for morphological parameters, like the length (Fig. 3H) and bending index (Fig. 3I), remained unchanged between fully- and semi-automated analyses, demonstrating that CiliaQ analysis can be used in a completely automated and high-speed analysis. However, the segmentation errors have to be carefully assessed beforehand, and a full-automated analysis can be only conducted with a low segmentation error. For example, a low signal-to-noise ratio in the cilia marker channel, high ciliary density, or a low imaging resolution can preclude fully-automated cilia detection.

### 2.3 2D versus 3D analysis

Although cilia in most tissues and cell culture stretch into all three dimensions, cilia have been mainly analyzed in 2D projections, which comes with a large bias of the results [32]. CiliaQ can be employed on both, 2D and 3D images, and thereby, allows to assess the bias of a 2D analysis. To this end, we analyzed a standard cilia data set in 2D and 3D. We acquired 3D confocal images of serum-deprived mIMCD-3 cells, and analyzed the images either directly (3D analysis) or performed a maximum intensity projection before analysis (2D analysis).

Furthermore, the analyzed mIMCD-3 cells were stained for two commonly used ciliary marker proteins, acetylated Tubulin (acTUB) and ARL13B [39] (Fig. 4A), as this allowed to test whether a 2D analysis alters the results obtained from different labeling methods equally, and to determine whether reconstructions from either marker differ. Thus, we also reconstructed the cilia based on each label.

**Figure 4:**
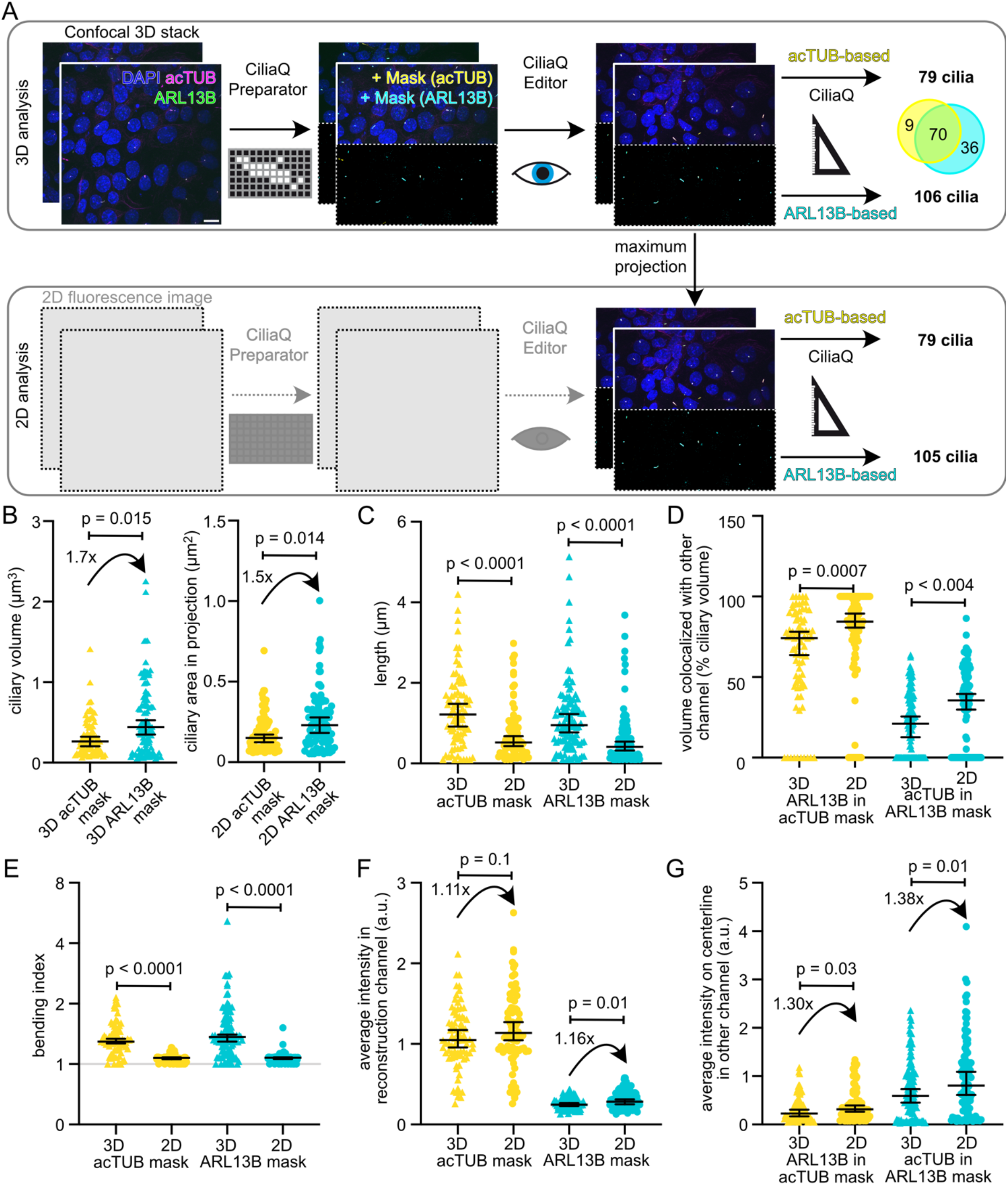
Comparing acetylated-Tubulin-to ARL13B-staining for ciliary reconstruction in 2D and 3D analysis. **(A)** Analysis workflow for the analyzed data set containing two confocal 2D stacks acquired from serum-starved IMCD-3 cells that were stained with acetylated Tubulin (acTUB) and ARL13B antibodies, both labeling cilia, and with DAPI to label nuclei. First, the image stacks were segmented using CiliaQ Preparator; both, the acTUB and the ARL13B channel, were segmented. Second, the acTUB and the ARL13B masks were independently corrected using CiliaQ Editor. Third, cilia were quantified in 3D with CiliaQ using either the acTUB or the ARL13B mask. The Venn diagram shows the overlap of cilia detection using both labels: 9 cilia were only detected with acTUB, 70 cilia were detected with acTUB and ARL13B, and 36 cilia were detected only with ARL13B. To mimic a 2D analysis paradigm of the identical cilia, CiliaQ was also applied to maximum intensity projections of the segmented and corrected stacks (bottom row). Scale bar: 10 µm. **(B-G)** CiliaQ analysis results for the four different types of analysis shown in A. Arrows in F and G indicate the fold change of 2D compared to 3D analysis. P-values indicate the test results of a Mann-Whitney test between the samples connected with a bar or an arrow.

Of note, we observed large differences in CiliaQ 3D analysis and results comparing the reconstruction from the acTUB and from the ARL13B staining. First, using the mask based on acTUB staining required more editing compared to the ARL13B mask because tubulin acetylation is not restricted to primary cilia only [39, 40]. Second, the ARL13B mask revealed more cilia than the acTUB mask (3D: 106 vs. 79, respectively), demonstrating that marker expression varies among cilia from mIMCD-3 cells. In 3D, 70 cilia were detected via either labeling method, 9 cilia were detected via acTUB labeling only, and 36 cilia were detected via ARL13B labeling only (Fig. 4A). Third, the acTUB-based 3D reconstruction revealed volumetrically smaller but longer cilia than the ARL13B-based 3D reconstruction (Fig. 4B-D), which indicates that ARL13B-based reconstructed cilia are thicker. The latter could be explained by the fact that tubulin acetylation occurs only in the core of the cilium [40], while ARL13B associates with the ciliary membrane [41]. The higher detection rate using ARL13B, rather than acTUB labeling, could relate to the fact that tubulin acetylation is reduced during cilia resorption [11]. Cilia positive for ARL13B but negative for acTUB could be in the progress of cilia resorption.

Taken together, the differences between the markers highlight that comparing cilia reconstructions based on different markers is not trivial. Furthermore, the use of the marker obviously has implications for protein localization-studies in cilia. Using cilia reconstruction based on a label for the core of the cilium (i.e. acTUB) reduces the detection of a colocalized protein that is localized in the ciliary membrane (i.e. ARL13B) (Fig. 4D). Thus, ARL13B appears also more suitable to study ciliary protein localization than acTUB.

In a 2D analysis of the same images, we identified 105 (ARL13B) and 79 (acTUB) cilia, revealing that the number of detected cilia was hardly affected by the 2D analysis (Fig. 4A). Furthermore, the trend of obtaining bigger cilia using ARL13B instead of acTUB for reconstruction was conserved in a 2D analysis (Fig 4B). Because the z component of the cilium is missing, the morphological output parameters, e.g. ciliary length (Fig. 4C) and bending index (Fig. 4E), were decreased in the 2D compared to the 3D analysis. Notably, the ciliary bending index, which relies on measuring the ciliary length and orientation, was largely different in 2D vs 3D (Fig. 4E): In 2D, the values approached 1 (average: 1.07 for acTUB and ARL13B), indicating no bending at all, whereas in 3D, a slight bending of about 1.35 (acTUB) and 1.50 (ARL13B) on average was detected. This shows that 2D analysis was not able to detect the fine difference in cilia bending between acTUB- and ARL13B-based cilia reconstructions. For intensity parameters, differences between 2D and 3D results were less significant (Fig. 4F-G). Taken together, this indicates that a 2D analysis of ciliary morphology is inaccurate and can preclude the detection of differences in morphology, whereas a 2D analysis of total ciliary intensity levels is less biased.

### 2.4 Applying CiliaQ to tissue images

To analyze cilia in tissues using CiliaQ, we recorded confocal 3D images of the developing spinal cord (Fig. 5) and telencephalon (Fig. 6) of zebrafish embryos. *In vivo* studies of cilia commonly employ transgenic models, expressing a ciliary protein fused to a fluorescent protein. Analogously, we analyzed *b-actin:arl13b-gfp* embryos that provide cilia labeling based on transgenic expression of ARL13B-GFP [29, 42]. Fixed embryos were additionally stained with an acTUB antibody as a second ciliary marker (Fig. 5A, C). To assess the effect of transgenic ARL13B-GFP expression, we segmented cilia from background based on the acTUB channel (Fig. 5B). Here, extensive editing was required because acTUB also labels structures other than cilia (e.g. neurons and axons) [40], and cilia at the ventral side of the spinal cord are very densely packed [43], resulting in many overlapping ciliary acTUB and ARL13B signals (Fig. 5A). Cilia that could not be clearly distinguished from each other where manually excluded.

**Figure 5:**
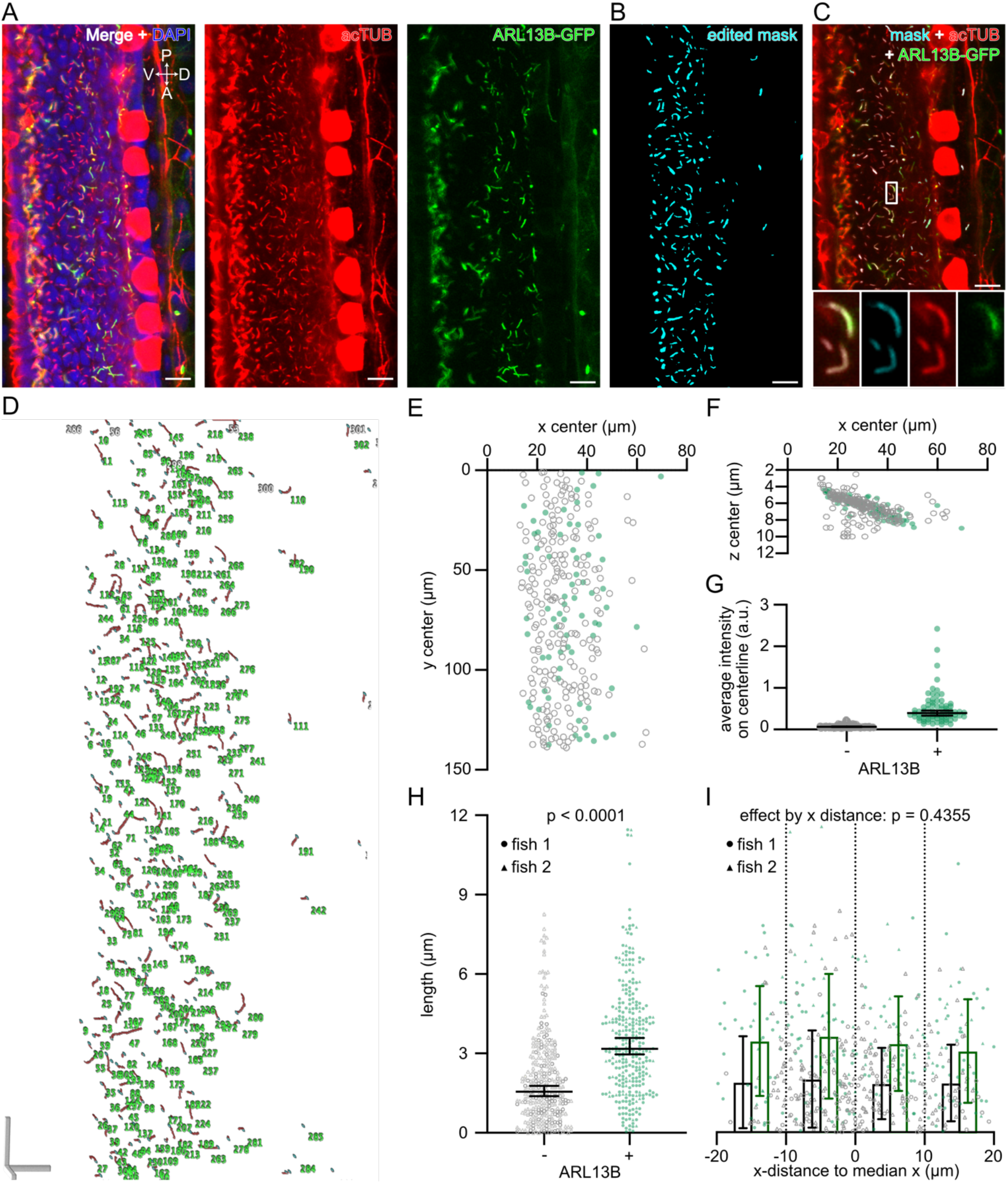
CiliaQ analysis of cilia in a tissue, i.e. the zebrafish spinal cord. **(A)** Exemplary slice from a confocal stack through the spinal cord of a zebrafish embryo (28 hours post fertilization), genetically expressing ARL13B-GFP some cells, stained with an acetylated Tubulin (acTUB) antibody to label cilia and DAPI to label nuclei. Left: all three labels. Middle: acTUB channel only. Right: ARL13B channel only. V = ventral, D = dorsal, P = posterior, A = anterior. **(B)** Corrected mask generated by segmentation and editing of the stack shown in A using CiliaQ Preparator and CiliaQ Editor, respectively. **(C)** Overlay of the corrected mask and the original channels. Magnified view of the position indicated with the white box is shown at the bottom for all channels merged (left) or the corrected mask (middle left), the acTUB channel (middle right), or the ARL13B channel (right) only. The images shown in A-C map the the same stack positions. **(D)** 3D visualization (skeleton representation output by CiliaQ), **(E-F)** coordinates and **(G)** ARL13B intensity of the cilia detected by CiliaQ analysis. **(H)** Length of the cilia detected by CiliaQ analysis. Results pooled from *n* = 2 fish. p-value for an unpaired, two-sided Mann-Whitney test indicated. **(I)** Data from H, replotted by distance of the cilia in x, to the median x position for each fish. p-value for the factor x-distance determined using a 2-way ANOVA (alpha = 0.05). Results for ARL13B-negative and -positive cilia were separated based on the CiliaQ parameter colocalized volume measured in the ARL13B channel: cilia with a colocalized volume above 0 were considered ARL13B-positive. Scale bars: 10 µm.

**Figure 6:**
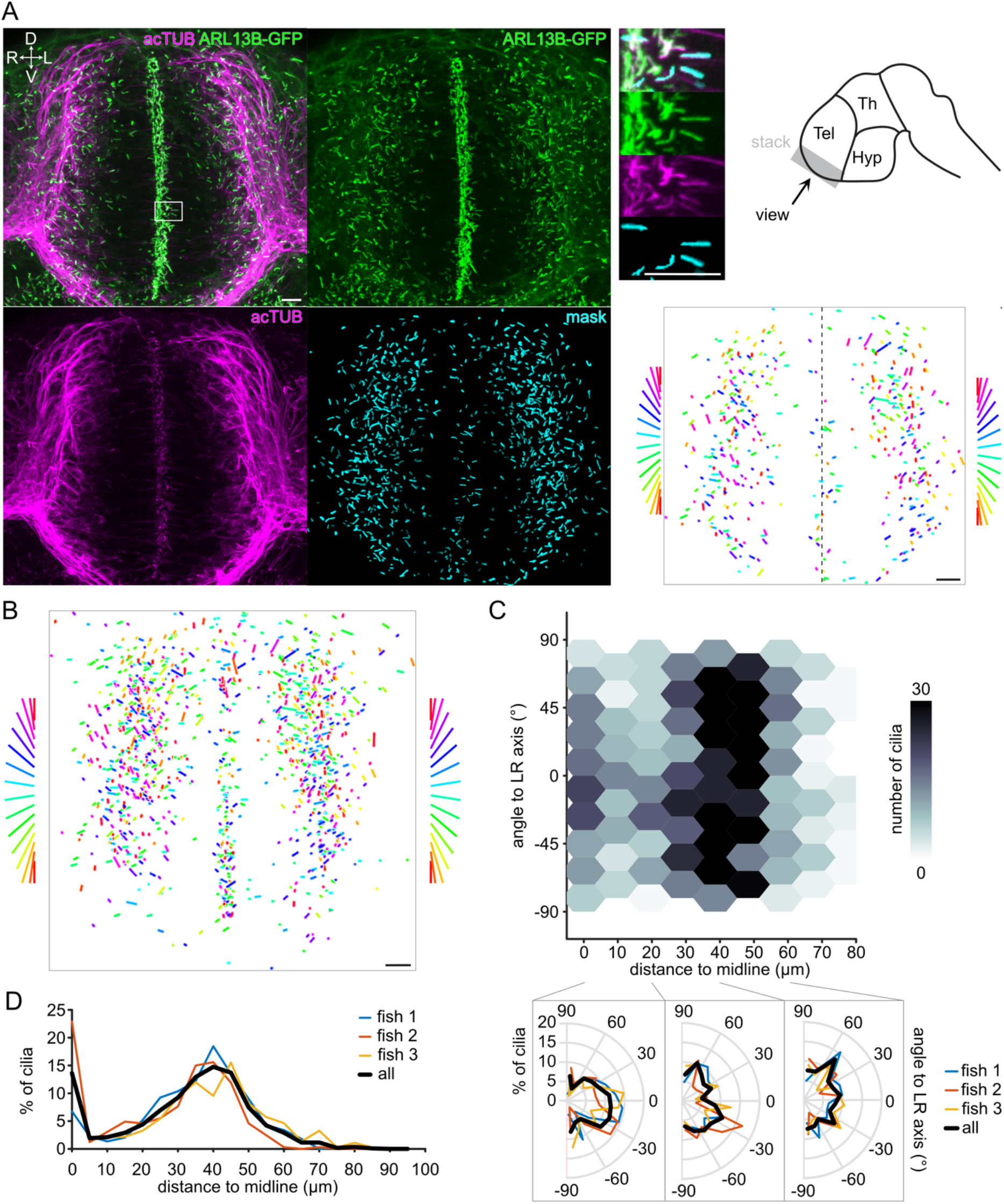
CiliaQ analysis of cilia orientation in the telencephalon of zebrafish embryos. **(A)** CiliaQ analysis of confocal stack through the telencephalon of a zebrafish embryo (28 hours post fertilization), genetically expressing ARL13B-GFP and stained with an acetylated Tubulin (acTUB) antibody. Left: Maximum intensity projection of the stack (overlay and individual channels) and of the mask derived from the ARL13B channel using CiliaQ Preparator and corrected using CiliaQ Editor. Top right: Magnified view of the maximum intensity projection and schematic drawing of the brain indicating from which region the stack (grey) shown on left has been recorded. Bottom right: plot of orientation vectors (only vectors included with a length >= 1 µm) derived by CiliaQ analysis of the mask stack. Orientation vectors are color-coded by direction, as indicated on the sides for the right and left side of telencephalon (separated by dashed line in image). D = dorsal, V = ventral, R = right, L = left. Tel = Telencephalon, Th = Thalamus, Hyp = Hypothalamus. **(B)** Overlay of the orientation vector derived by CiliaQ analysis of image stacks from *n* = 3 embryos. **(C)** Replotting orientation vectors shown in B (only x and y components considered) as a function of the distance to the midline (as indicated in A), representing the brain ventricle. Bottom: circular histograms of the orientation vectors in the ranges of 0-20 µm, 20-40 µm, and 40 – 60 µm distance to the midline. Results indicated for individual fish and for all fish pooled. Overlay of the corrected mask and the original channels. Magnified view of the position indicated with the white box is shown at the bottom for all channels merged (left) or the corrected mask (middle left), the acTUB channel (middle right), or the ARL13B channel (right) only. The images shown in A-C map the same stack positions. Number of cilia in percent of all detected cilia as a function of the distance to the midline for the *n* = 3 fish shown in B and C. Data are shown by individual fish and for all fish pooled into one data set. Scale bars: 10 µm.

It has been reported that overexpression of a ciliary proteins increases cilia length [34, 44-46]. To test whether this is also the case in the *b-actin:arl13b-gfp* embryonic spinal cord, we compared ARL13B-GFP-positive and negative cilia (Fig. 5C). CiliaQ reconstructed 293 cilia in one image (Fig. 5D). ARL13B-GFP-positive or negative cilia were identified based on the parameter ‘colocalized length’ in the GFP channel: Cilia with a colocalized length above 0 were considered ARL13B-GFP-positive, while cilia with a colocalized length of 0 were considered ARL13B-GFP-negative. ARL13B-GFP-positive and -negative cilia were distributed across the entire field of view (Fig. 5E-F). ARL13B-GFP-positive cilia showed a significantly higher average intensity on centerline (Fig. 5G) and were generally longer than ARL13B-GFP-negative cilia (Fig. 5H). We did not detect any difference in cilia length on the dorsal-ventral axis (Fig. 5I). However, motile cilia at the ventral side of the spinal cord, at the level of the central canal [43], are not well represented, as we excluded ciliary reconstructions of overlapping adjacent cilia.

We next investigated cilia in the telencephalon. Because not all cilia in the telencephalon were clearly acTUB-positive, we analyzed cilia based on the Arl13B-GFP channel (Fig. 6A). Again, we excluded cilia, this time at the ventricle, where we could not clearly separate adjacent cilia belonging to highly polarized neuroprogenitors [34, 47-50]. CiliaQ allowed to precisely determine the orientation of the cilia in relation to the ventricle (midline) (Fig. 6A, bottom right). Most cilia close to the midline, belonging to delaminating neurons [51, 52], aligned with the left-right axis, while neuronal cilia in the lateral part of the developing telencephalon pointed to all directions. We further confirmed this finding in two more embryos (Fig. 6B) and represented the orientation of cilia in different regions using polar histograms (Fig. 6C). Of note, we also uncovered that most cilia in the telencephalon lie either close to the ventricle or at a distance of around 40 µm to the ventricle (Fig. 6D).

We conclude that CiliaQ can also handle large 3D images, such as images of whole organs or tissues, which allows to gain new insights into orientation, morphology, and protein composition of cilia.

### 2.5 Error assessment in fully-automated analysis of tissue images

Finally, we quantitatively assessed the errors of a fully-automated CiliaQ analysis using a tissue image, i.e., of the developing spinal cord in a zebrafish embryo (Fig. 7A). Here, many cilia are closely adjacent, whereby the ciliary masks overlap, even in 3D confocal-microscopy images.

**Figure 7:**
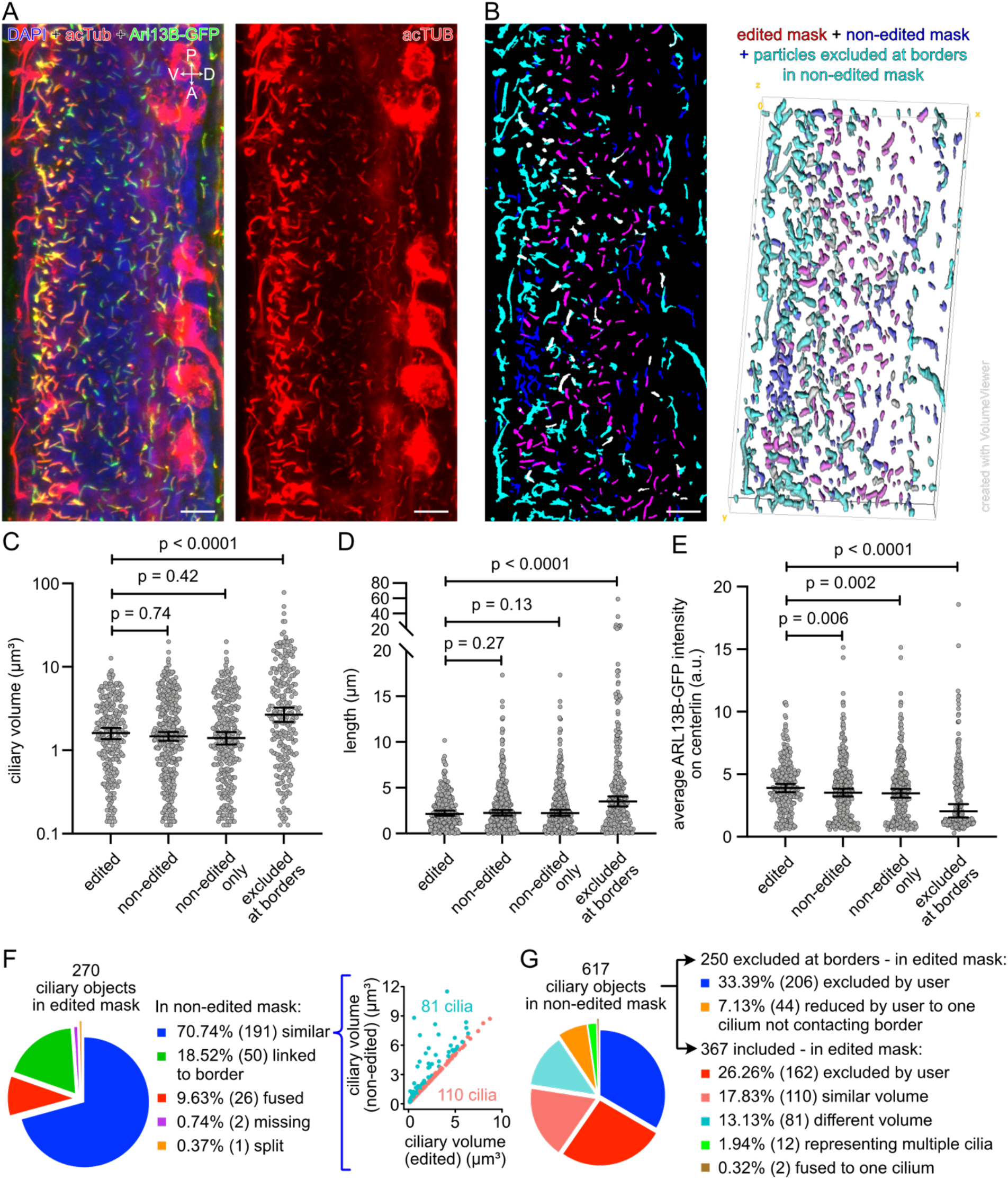
Accuracy of automated CiliaQ analysis using an image of the spinal cord in a zebrafish embryo. **(A)** Maximum projection of a confocal stack through the spinal cord of a zebrafish embryo (28 hours post fertilization), genetically expressing ARL13B-GFP in some cells, stained with an acetylated Tubulin (acTUB) antibody to label cilia and DAPI to label nuclei. Left: all three labels. Right: acTUB channel only. V = ventral, D = dorsal, P = posterior, A = anterior. **(B)** Overlay of (1) the edited mask (red), generated by using CiliaQ Preparator, CiliaQ Editor for manual correction of the segmentations, and CiliaQ for excluding particles touching borders or not exceeding a size threshold (10 voxel), of (2) the non-edited mask (blue), generated as the edited mask but without manual corrections, and of (3) particles that were excluded from the non-edited mask by CiliaQ due to touching the borders (cyan). Left: Maximum projection of the mask stack. Right: 3D rendering of the mask stack. Objects in magenta are present in both, the edited and non-edited mask (mixing color of blue and red). Objects in white represent cilia that were manually included but excluded from the non-edited mask because of fusion to particles that were connected to the borders of the image (mixing color of cyan and red). **(C-E)** CiliaQ output parameters determined based on analyzing the masks shown in B. Results are plotted by different groups of cilia: (1) ciliary objects detected by analyzing the edited mask, (2) ciliary objects detected by analyzing the non-edited mask (excluding cilia at the border of the image in CiliaQ), (3) ciliary objects that were detected only in the non-edited mask, (4) ciliary objects excluded from the non-edited mask in CiliaQ because of touching the borders of the image. p-values for Kolmogorov-Smirnov tests, which compare the cumulative distributions, are indicated. **(F-G)** Comparative analysis of the ciliary objects reconstructed based on the edited and based on the non-edited mask. (F) Left: Appearance of ciliary objects reconstructed from the edited mask in the non-edited mask. Right: Correlating volumes determined based on the edited to volumes determined based on the non-edited mask. Cyan: cilia with a different volume in non-edited and edited mask (volumetric difference > 1.1x). Rose: cilia with similar volume in non-edited and edited mask (volumetric difference < 1.1x). (G) Appearance of ciliary objects reconstructed from the non-edited mask in the edited mask. Scale bars: 10 µm. Cilia objects from the edited mask are also presented in Fig. 5G-H as fish 2.

We compared the results from analyzing a manually-corrected edited mask with a non-edited mask without any manual correction (Fig. 7B). Overall, the results of the morphological parameters like the ciliary volume (Fig. 7C) and ciliary length (Fig. 7D) did not differ between the two masks. In either case, the standard CiliaQ workflow, as applied here, automatically excludes ciliary objects touching the image borders. When analyzing specifically the particles that were excluded from the non-edited mask because they touched the image borders, we observed a significantly increased ciliary volume and length. We related this to the observation that in the non-edited mask, without user-dependent corrections, multiple closely adjacent cilia were fused to one particle and such agglomerated particles are more likely to touch the image borders because they span a larger volume than particles representing one cilium only. In conclusion, excluding particles touching the border of the image allowed by large to exclude incorrectly detected ciliary objects that represent multiple overlapping cilia in the fully-automated analysis.

In contrast, intensity parameters differed between the non-edited and edited mask (Fig. 7E). We observed a consistent reduction in intensity between the edited and non-edited mask. This indicates that an improper mask, detecting ciliary objects that fuse multiple cilia, resulted in a dilution of measured intensity signals, as not all cilia were ARL13B-GFP positive. We sought to quantify this in detail and individually compared 3D objects in both masks. For each particle in the edited mask, we linked particles in the non-edited mask that overlapped by at least one voxel: 191 of 270 cilia (70.7%) overlapped with only one particle in the non-edited mask, from which 110 (40.7% of all 270 edited cilia) revealed a similar volume in the non-edited mask (Fig. 7F). 81 cilia (30% of all 270 edited cilia) showed a more than 10% higher volume in the non-edited compared to the edited mask, indicating fusion to adjacent objects that were removed by the user. 26 cilia (9.6%) were confirmed to be fused into less ciliary objects in the non-edited mask, while only one cilium was split into two ciliary objects in the non-edited mask (Fig. 7F). We also tested how many cilia were correctly detected from the non-edited mask by linking particles from the edited mask to particles in the non-edited mask based on overlap (Fig.7G). 250 of 617 (40.5%) of all ciliary objects in the non-edited mask were excluded as they touched the image borders, while 44 of them overlapped with cilia that were revealed by the user in the edited mask. From the remaining 367 ciliary objects, the majority (162 ciliary objects) could not be corrected by the user, as it was not possible to decide whether they represent overlapping cilia or were non-ciliary structures. 30% of the included ciliary objects (110) were confirmed as correct, as they overlaped with the edited mask. Thus, we conclude for the presented data set that errors caused by overlapping cilia appear more frequent than errors caused by inhomogeneous labeling of the cilium.

Last but not least, it is important to consider that the errors described here cannot be directly inferred for other data-sets. The detection of ciliary objects and the concomitant diverse types of errors are data-set-specific and depend on the type of cell/tissue analyzed, the ciliary labeling technique [39], and the imaging set-up. For example, besides errors from fusion and splitting cilia into ciliary objects, errors from detecting staining artefacts or non-ciliary structures labeled by a ciliary marker, e.g., the spindle apparatus [40], may be dominant in other data-sets. To establish an automated CiliaQ analysis-pipeline, it is important to assess and consider dataset-specific error rates. In the present data set, we faced the disadvantage that the diversity of cilia morphology was high, as the data set contained two classes of cilia with largely different length and volume. For populations where the variance in ciliary morphology parameters is low, it might be possible to automatically detect and exclude incorrect ciliary objects that represent multiple overlapping cilia by applying thresholds for morphological parameters.

## 3 Discussion

We present an open-source software for analyzing ciliary morphology and fluorescence intensity in 2D, 3D, and, if applicable, time-lapse 2D and 3D images of tissue cells *in vivo, in situ*, and in tissue culture. The ImageJ plugins, the underlying source-code, and a user guide are freely-accessible through the online GitHub repository (https://github.com/hansenjn/CiliaQ). The application of CiliaQ does not require any coding knowledge – CiliaQ integrates as plugins into the broadly used, open-source, freely-available image analysis software ImageJ [53, 54]. We equipped CiliaQ with a multi-task-management system to use CiliaQ for automated analysis of large data sets. We envision a broad applicability of CiliaQ in cilia research.

Other tools and approaches to automatically detect and reconstruct cilia of tissue cells in digital images have been described, starting in the eighties [31-33, 55]. However, these studies are either limited to the characterization of individual parameters, e.g. only the ciliary centerline in 2D [55], the ciliary length in 3D [32, 33], the length and frequency (% of ciliated cells) in 2D [31], the ciliary orientation and distribution [56], or to characterize individual cilia per image [24, 57]. Notably, the method by Ferreira et al. [56] requires manual optimization of the imaging setup and relies on inferring ciliary orientation from imaging artefacts, thus does not allow a trivial, precise reconstruction of ciliary morphology parameters like the ciliary length or bending. Furthermore, the semi-automated “Simple Neurite Tracer” [58] has been transferred to cilia analysis [59]. However, this method requires manual tracing of the cilium, which can be sensitive to user bias and provides low throughput.

In contrast, CiliaQ provides a comprehensive list of different parameters for the morphology, orientation, bending, and protein content of many cilia in 2D images, 3D images, and time-lapse 2D or 3D images. CiliaQ automatically obtains ciliary 3D regions using image segmentation, for which we implemented a large variety of segmentation approaches into CiliaQ: a common intensity thresholding, 3D hysteresis thresholding, or a method based on the Canny edge detection [35] (Canny3D). CiliaQ allows users to flexibly select the best suited segmentation approach for their data set. For example, we show that a common intensity thresholding method more precisely retrieves the ciliary structure for homogeneously stained cilia, while Canny3D is better suited for unevenly labeled cilia as it involves edge detection and generates larger ciliary reconstructions. Taken together, CiliaQ provides a very flexible whilst comprehensive analysis method of cilia.

Multiple methods have been proposed to study the ciliary length. For example, cilia length has been determined based on the Pythagorean theorem [32]. However, this only applies to perfectly straight cilia, while high cilia bending will bias the analysis. Similarly to a reported method for precisely estimating ciliary length in 3D [33], CiliaQ employs automated scaling, blurring, and skeletonization of the ciliary 3D region to obtain good 3D length estimations also from low signal-to-noise-ratio images.

Here, we present different applications of CiliaQ - from a simple analysis of the ciliary SMO localization in cultured cells to studying ciliary orientation in the developing zebrafish telencephalon. We highlight that the comprehensive analysis of CiliaQ is very fast and completely user-independent for individual cells. For such standard applications of CiliaQ, i.e. stained cells in culture, we show that CiliaQ provides a precise, fully-automatized 3D (and also time-lapse 3D [34]) analysis, allowing its application even in high-throughput image analysis. Moreover, we show that a 2D analysis is largely biased, especially for morphological parameters like cilia length and bending, while fluorescence parameters for the total protein content are less affected, which is in line with previous comparisons of 2D and 3D analysis [32, 33]. However, 3D analysis has been challenging for a standard biology lab, whereby many labs preferred to perform a manual 2D analysis. Hence, CiliaQ was developed to handle the full range of simple 2D imaging settings up to sophisticated time-lapse 3D spinning-disk microscopy [34].

We also demonstrate detection of cilia by CiliaQ with different labelling methods, i.e. antibody staining for two widely established ciliary markers, ARL13B and acetylated-Tubulin (acTUB) [39], and overexpression of a ciliary protein labeled with a fluorescent protein [42]. We demonstrate that results are dependent on the choice of ciliary marker, i.e. morphology and intensity parameters differed between ARL13B- and acTUB-based reconstructions. Notably, we show that an acTUB-based reconstruction incompletely captures ciliary localization of a membrane protein (ARL13B) and that some cilia of mIMCD-3 cells were ARL13B positive, but not acTUB positive and vice-versa. A previous study comparing different ciliary markers reported that the fixation method influences ciliary labeling [39]. Thus, it is highly recommended to carefully select the ciliary marker and keep the fixation method and the used ciliary marker consistent within a data set. In addition, we observe that cilia labeled by overexpressing a cilium-localized fluorescent protein appeared longer than antibody-labeled cilia. In line with previous reports [34, 44, 45], this highlights that the amount of overexpressed ciliary protein needs to be carefully titrated to avoid biasing ciliary morphology.

For images where individual neighboring cilia are indistinguishable (i.e. in tissues with multi-ciliated cells [60]) or where cilia have been labeled with a marker that also labels structures other than cilia (i.e. spindle apparati in an acTUB staining [39, 40]), CiliaQ still requires the user to correct the segmentation of cilia from background and to exclude other structures than cilia from the analysis. To this end, we integrated CiliaQ Editor into the CiliaQ workflow. CiliaQ Editor renders the CiliaQ workflow into a user-assisted but still mostly automatized pipeline, allowing to facilitate and standardize the analysis of such complicated images, which so far has been impossible or relied on purely manual analysis.

We demonstrate that CiliaQ can be used to study and map cilia in tissues like the spinal cord and developing telencephalon of the zebrafish embryo. Reconstruction of cilia in these tissues was more challenging and thus, required additional manual editing. The main limitations were associated with a high density of cilia at the midline or central canal limiting an error-free segmentation, the lack of highly specific ciliary markers resulting in an elevated background staining, and the influence of the transgenic expression of a ciliary marker (e.g. ARL13B-GFP) on ciliary length. Despite these challenges, our analysis allows to gain new insights into cilia and developmental biology. In particular, we observed that cilia are spatially organized and polarized in the developing zebrafish telencephalon, but not in the spinal cord. In the developing telencephalon, we uncovered that neuronal progenitors extend their cilia towards fluid filled cavities [29], similarly to mammalians and chicken embryos [47-50]. Moreover, we observed that during neuronal delamination, cilia are oriented along the left-right axis, and that this orientation vanishes when the neurons are differentiated and start expressing acTUB.

As an open-source project, we envision CiliaQ, with its GitHub repository, as a lively platform to connect researchers to further automatize and optimize cilia analysis, for example for densely ciliated tissues. Here, new approaches to separate cilia in densely ciliated tissues and distinguish cilia from other labeled structures are required. CiliaQ can be used as a starting point to develop new and more specialized analysis approaches, which can also be included into CiliaQ later. Because CiliaQ is broadly applicable to many types of experiments, freely available, and open-source, introducing new features for a specific analysis will make them directly available to a much broader community.

## 4 Materials and Methods

### 4.1 Cell lines and cell culture

mIMCD-3 cells (mouse Inner Medullary Collecting Duct 3 cells) were obtained and authenticated from American Type Culture Collection (ATCC, CRL-2123). mIMCD-3 were maintained in DMEM/F12 (1:1) medium, supplemented with GlutaMax (both: Gibco(tm), Thermo Fisher) and 10 % FCS at 37°C and 5 % CO_2_.

Mouse Embryonic Fibroblasts (MEFs) were isolated from E13.5 wildtype embryos and immortalized according to the method of Todaro and Green [61]. MEF cells were maintained in DMEM medium, supplemented with 1% Penicillin-Streptomycin-Glutamin, 1% sodium pyruvate (all: Gibco(tm), Thermo Fisher), and 10% FCS at 37°C and 5% CO_2_.

All cells have been tested and are free from mycoplasma and other microorganisms.

### 4.2 Immunocytochemistry of cultured cells

Immunocytochemistry was performed according to standard protocols. Cells were seeded on poly-L-lysine (PLL, 0.1 mg/ml, Sigma Aldrich)-coated 13 mm glass coverslips (VWR) in a 4-well dish (VWR). The next day, the medium was replaced with starvation medium (0.5% FCS) to induce ciliogenesis. After 24 hours of starvation, MEF cells were stimulated for 24 hours with Smoothened agonist (SAG, 1 µM, Sigma-Aldrich) or the solvent as control and next washed with PBS, fixed with 4 % paraformaldehyde (Alfa Aesar, Thermo Fisher Scientific) for 10 min at room temperature and washed again with PBS. mIMCD-3 cells were fixed 24 or 48 hours after starvation. After washing with PBS, cells were blocked with CT (0.5% Triton X-100 (Sigma Aldrich) and 5% ChemiBLOCKER (Merck Millipore) in 0.1 M NaP, pH 7.0) for 30 minutes at room temperature. Primary and secondary antibodies were diluted in CT and incubated for 60 min at room temperature. Coverslips were mounted with one drop of Aqua-Poly/Mount (Tebu-Bio). The following antibodies were used: mouse anti-acetylated-Tubulin (1:600, Sigma Aldrich, T6793), rabbit anti-ARL13B (1:500, Proteintech, 17711-1-AP), mouse anti-ARL13B (Abcam, ab136648, 1:500), rabbit anti-Smo (1:500, Anderson lab [62]), mouse anti-gamma-Tubulin (Sigma Aldrich, T6557, 1:2000), donkey anti-mouse-Cy3 (1:1000, Dianova, 715-165-151), donkey anti-mouse-Cy5 (1:500, Dianova, 715-175-151), goat anti-rabbit-Alexa488 (1:500, Life Technologies, A11034). As a DNA counterstain, DAPI was used (4’,6-Diamidino-2-Phenylindole, Dihydrochloride, 1:10 000, Invitrogen).

### 4.3 Confocal microscopy of cultured cells

Confocal z-stacks (step size 0.4-0.5 µm, 60x objective) were recorded with a confocal microscope (Eclipse Ti, Nikon). All depicted images show a maximum projection of a z-stack unless differently stated in the figure legend.

### 4.4 Zebrafish as an experimental model

The animal facilities and maintenance of the zebrafish, *Danio rerio*, were approved by the Norwegian Food Safety Authority (NFSA, 19/175222). Fishes were kept in 3.5 l tanks in a Techniplast Zebtech Multilinking system at 28 °C, pH 7 and 700 mSiemens, at a 14:10 h light/dark cycle. Fish were fed dry food (ZEBRAFEED; SPAROS I&D Nutrition in Aquaculture) two times/day and *Artemia nauplii* once a day (Grade0, platinum Label, Argent Laboratories, Redmond, USA). Embryos were maintained in egg water (1.2 g marine salt and 0.1% methylene blue in 20 l RO water) from fertilization on. All procedures were performed on zebrafish embryos in accordance with the directive 2010/63/EU of the European Parliament and the Council of the European Union and the Norwegian Food Safety Authorities. For experiments the following zebrafish lines was used: *b-actin:arl13b-gfp* [42].

### 4.5 Immunocytochemistry of zebrafish embryos

Euthanized embryos were fixed in a solution containing 4% PFA in PBS for at least 2h at room temperature. Following fixation, samples were washed with 0.3% PBSTx (3 x 5 min), permeabilized with acetone (100% acetone, 10 min incubation at −20°C), washed with 0.3% PBSTx (3 x 10 min) and blocked in 0.1% BSA/0.3% PBSTx for 2 h. Embryos were incubated with acetylated tubulin antibody (clone 6-11B-1, MABT868, Sigma-Aldrich, 1:1000) overnight at 4 °C. On the next day samples were washed (0.3%PBSTx, 3 x 1 h) and incubated with the secondary antibody (Alexa-labelled goat anti-mouse 555 plus, A32727, Thermo Fisher Scientific, 1:1000), an anti-GFP antibody conjugated with Alexa 488 (A-21311, Thermo Fisher Scientific, 1:1000) and DAPI (D1306, Thermo Fisher Scientific, 1:1000) overnight at 4 °C. The next day samples were washed (0.3% PBSTx, 3 x 1 h) and transferred to a series of increasing glycerol concentrations (25%, 50% and 75%).

### 4.6 Confocal microscopy of zebrafish embryos

Stained embryos were mounted in 75% glycerol at 4 °C and imaged using a Zeiss Examiner Z1 confocal microscope with a 20x plan NA 0.8 objective. All depicted images show a maximum projection of a z-stack unless differently stated in the figure legend.

### 4.7 Image analysis

For all presented data sets, we summarize the Image specifications, CiliaQ Preparator preferences, and CiliaQ preferences in Table 2.

**Table 2:**
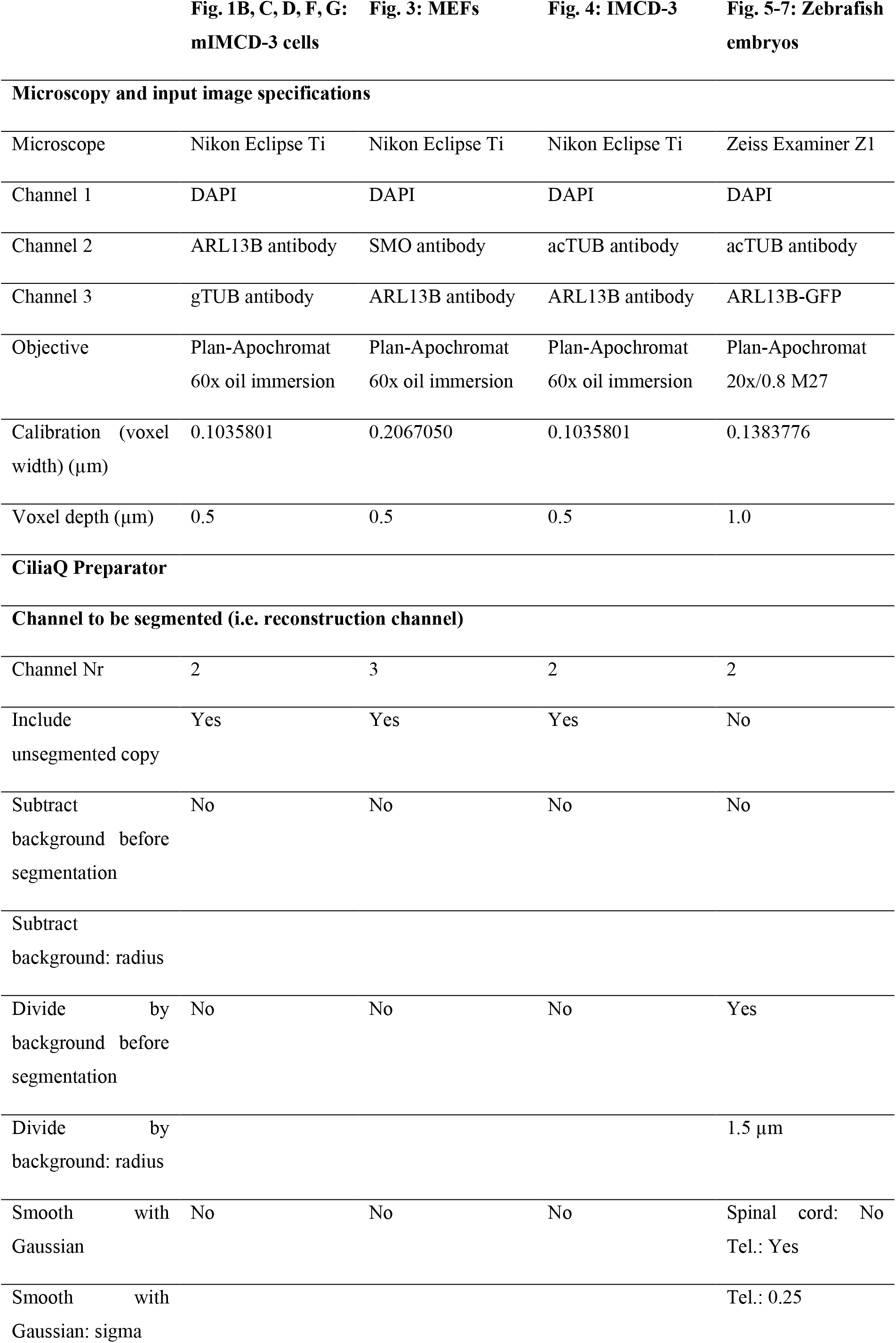
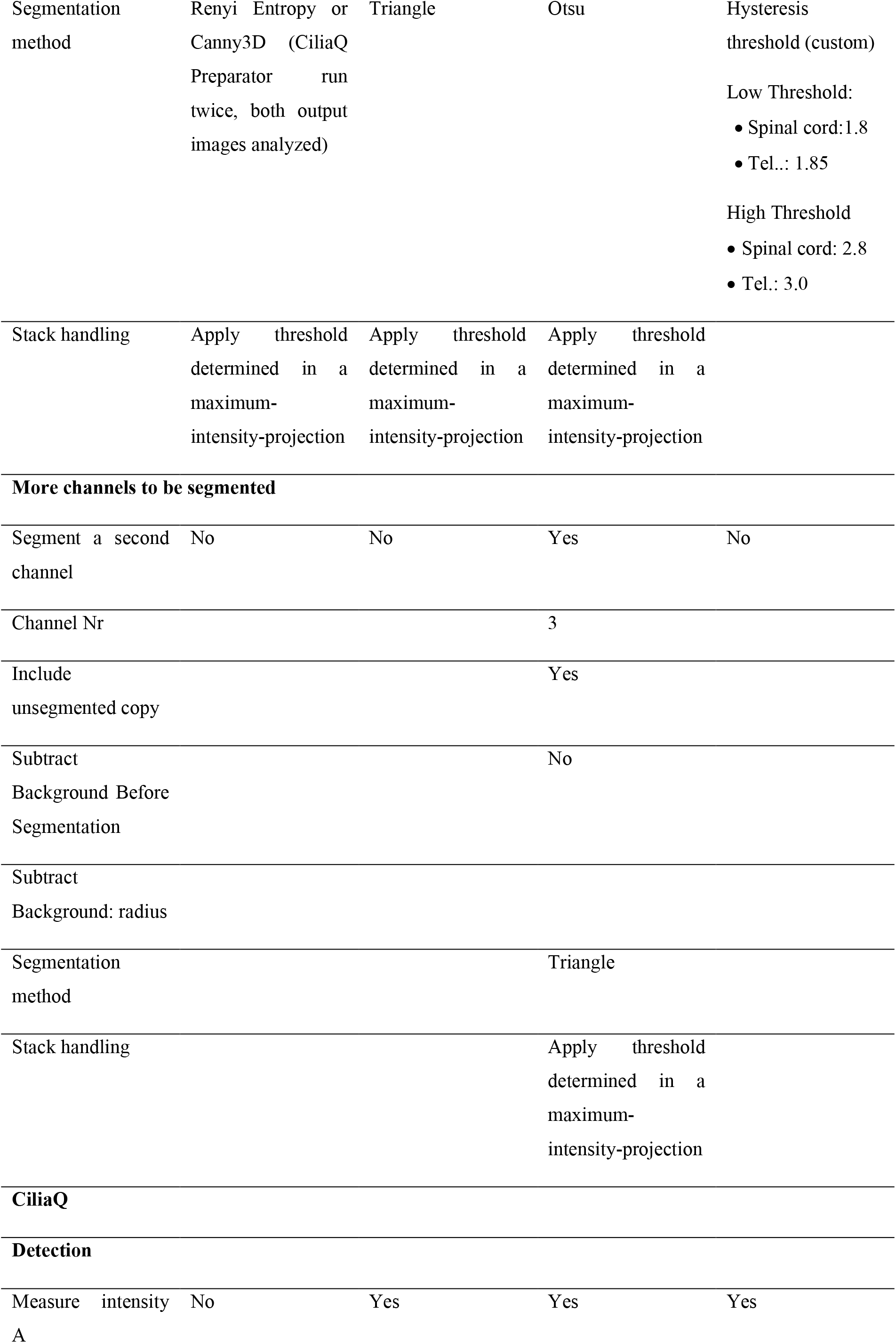
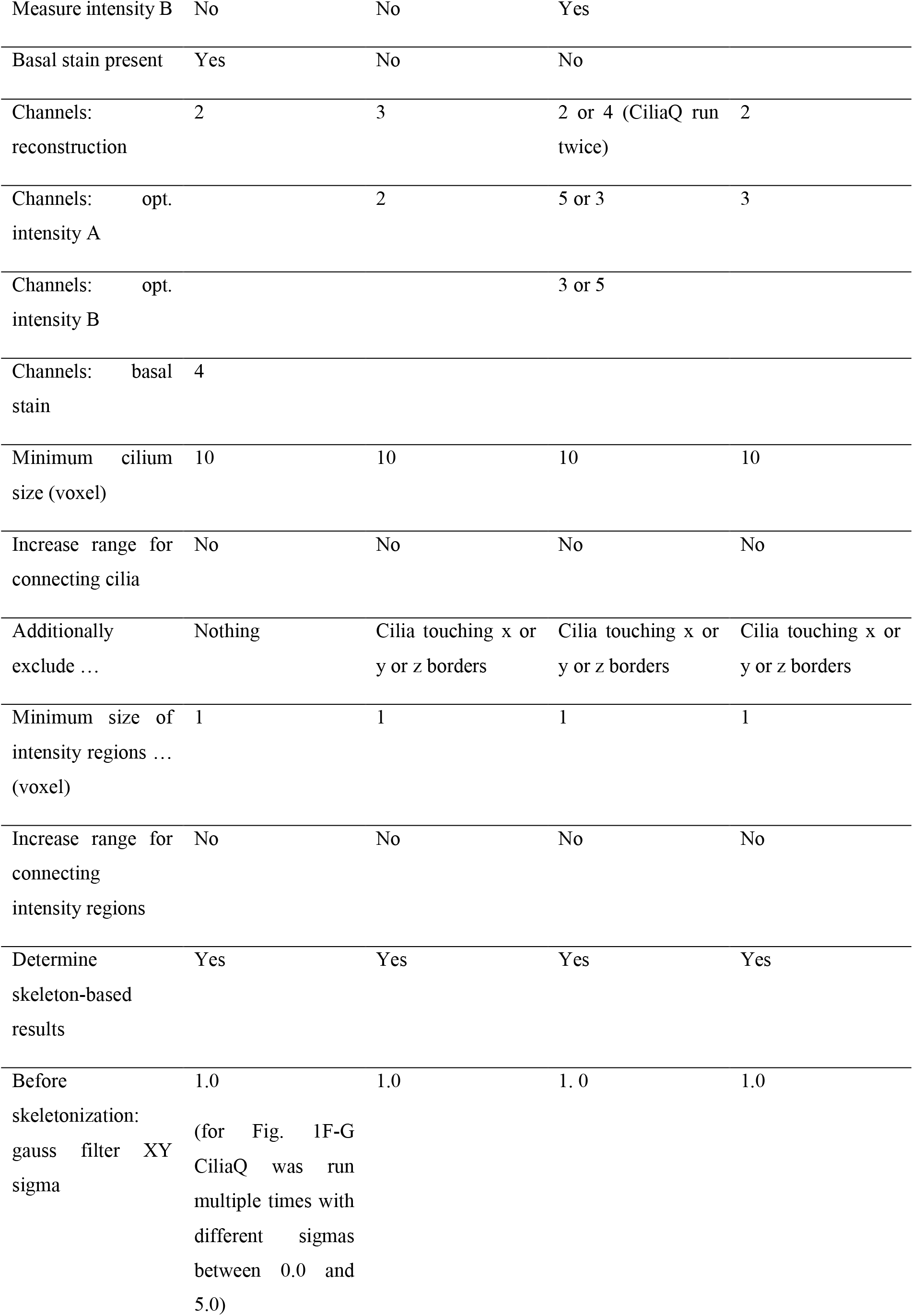
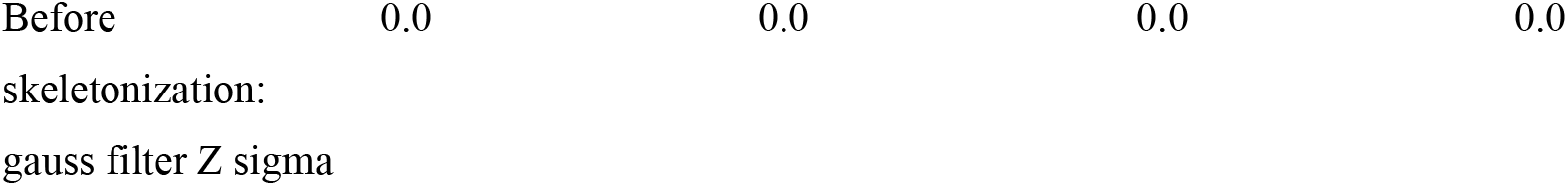
CiliaQ settings for presented data sets. Settings not applicable for the analysis result (e.g. visualization settings, settings left to default parameters) were not included in this table. Empty cells indicate that the entered value in the settings dialog is obsolete. Abbreviations: mIMCD-3 (mouse inner medullary collecting duct 3), MEFs (mouse embryonic fibroblasts), acTUB (acetylated Tubulin), gTUB (gamma Tubulin), tel (Telencephalon), voxel (represents a pixel in a 3D image).

### 4.8 Technical description of the CiliaQ plugins

Here we technically explain the technology underlying the CiliaQ plugins (CiliaQ Preparator v0.1.0, CiliaQ Editor v0.0.3, CiliaQ v0.1.4). We envision CiliaQ as a lively platform for cilia analysis that can be modified and extended in the future to enhance performance and applicability for cilia analysis. Thus, to obtain up-to-date information on the functioning, a user guide, and the current source code, please visit the GitHub page (https://github.com/hansenjn/CiliaQ).

#### 4.8.1 CiliaQ Preparator (version v0.1.0)

CiliaQ Preparator contains a “multi-task manager”, which allows to define an unlimited list of files for processing. The listed files, upon launching the processing, will then be processed one after the other, all under the identical settings.

Processing is completely automated and contains preprocessing and segmentation steps, which can be independently selected for each channel in the image when the plugin is launched (a list of preferences for the data sets presented here is shown in Table 2).

For preprocessing and segmentation of a channel, a copy of the channel to-be-processed is generated. Before segmentation, the channel’s images can be preprocessed by the following functions to reduce background and noise: (1) Background subtraction, using the ImageJ function Subtract Background (rolling ball) with a radius specified by the user. (2) Divide by background. The image is divided by a copy of the image that is processed with a Gaussian blur whose sigma is specified by the user. (3) Apply a Gaussian blur with a user-defined sigma.

For segmentation the following options are available: (1) ImageJ’s core auto-threshold methods, (2) the Canny3D method, (3) a Hysteresis threshold using the implemented intensity thresholds or custom thresholds, or (4) a custom threshold value can be selected. The individual methods are described in detail below.

Ultimately, each segmentation method will provide a list of cilia voxels (a voxel is the 3D representation of a pixel). This information is transferred to the original, unchanged channel in the input image to segment this channel. For this step, CiliaQ Preparator offers two variants:

(1) a binary image is generated, where the intensities of all detected cilia voxels are set to the highest intensity value (e.g. 255 a.u. for an 8-bit, 65535 a.u. for a 16-bit image) and the intensities of all other voxels are set to 0, or (2) a background-removed image is generated, where the intensities of all background voxels are set to 0 and the intensities of all cilia voxels remain unchanged. The latter approach features the advantage that cilia intensities are preserved and can be later quantified.

If selected by the user, the channel is duplicated before segmentation, so that the output image will contain the raw channel and the segmented channel.

The output image is automatically saved to the file location of the input image, under the name of the input image extended with the suffix “_CQP”. Additionally, a text file is saved, that stores all information on the preferences selected by the user. The log file can also be loaded into CiliaQ Preparator to reproduce the analysis or to apply identical preferences for additional images.

##### 4.8.1.1 Segmentation with an intensity threshold

The intensity threshold is automatically determined with the threshold methods implemented in ImageJ (IJ_IsoData, Huang, Intermodes, IsoData, Li, MaxEntropy, Mean, MinError, Minimum, Moments, Otsu, Percentile, RenyiEntropy, Shanbhag, Triangle, Yen) [53, 54]. Alternatively, a custom threshold value can be specified by the user. Next the image is segmented into cilia and background voxels based on the threshold: voxels with an intensity below the threshold are considered background voxels while voxels with an intensity equal to or above the threshold are considered foreground voxels (also referred to as “cilia voxels”).

For 3D images, the automatic intensity threshold method can be employed either on the histogram of a maximum intensity projection of the image stack or on the histogram of the whole image stack. For cilia, the histogram of a maximum intensity projection is commonly better suited than the stack histogram because in a standard 3D image of cilia, cilia make up only a small portion of the whole image’s volume and thus, are underrepresented in the stack’s intensity histogram. In the intensity histogram of a maximum projection the ratio of cilia voxels to background voxels is increased and thus, intensity populations of cilia and background voxels may be better distinguishable.

For time-lapse images, CiliaQ Preparator allows to determine one threshold for the whole time series or to determine individual thresholds for each time step. If the latter is selected, each individual time step will be segmented based on its respective individual threshold.

##### 4.8.1.2 Segmentation with a hysteresis threshold

For hysteresis thresholding, CiliaQ Preparator calls the function “3D hysteresis thresholding” implemented in the “3D ImageJ Suite” [36], an open-source software extension for ImageJ. Hysteresis thresholding requires to specify two intensity thresholds, one low threshold and one high threshold. For thresholding, the image is segmented into three groups of voxels: (1) voxels with an intensity below the low threshold, (2) voxels with an intensity equal to or above the high threshold, (3) voxels with an intensity equal to or above the low threshold and below the high threshold. Based on these three groups, the image is further segmented into fore- and background as follows: voxels from group (1) will be considered background; voxels from group (2) will be considered foreground (“cilia voxels”); voxels from group (3) will be considered foreground if they connect to voxels from group (2) and considered background if they do not.

CiliaQ Preparator allows to either automatically determine the low and high thresholds by selecting one of the threshold methods implemented in ImageJ for either threshold. Alternatively, custom threshold values can be specified by the user. CiliaQ Preparator allows to determine the threshold in the histogram of the maximum projection of the image or the histogram of the whole image stack (see above). Furthermore, CiliaQ Preparator allows to determine the threshold for the whole time series or to determine individual thresholds for each time step (see above).

##### 4.8.1.3 Segmentation with Canny3D

The Canny3D method that we developed represents a 3D adaption of Canny edge detection [35]. Canny3D employs four consecutive steps: (1) the image is smoothed with a 2D Gaussian kernel using the “Gaussian Blur” function implemented in ImageJ [53, 54], (2) edges are detected with a 3D Sobel kernel, (3) a 3D hysteresis threshold is applied, and (4) holes encapsulated in all three dimensions are filled. In step (1), a 2D and not a 3D kernel is applied, because in common fluorescence (confocal) microscopy the z-dimension is less resolved than the xy-dimensions and thus anyways blurred. In steps (2) to (4), functions from the ‘3D ImageJ Suite’ [36] are used. The user specifies the sigma of the Gaussian blur for step (1), the alpha of the 3D Sobel kernel for step (2), and the options to calculate low and high thresholds for step (3). Options for the latter are also explained in the previous section.

#### 4.8.2 CiliaQ Editor (version v0.0.3)

CiliaQ Editor opens the output image from CiliaQ Preparator and allows to remove voxels from or add voxels to the cilia voxels in the segmented channel. The segmented channel is specified by the user. The user is then requested to draw regions-of-interest (ROIs) around the regions in the image stack that shall be added or removed. Each applied ROI is saved, allowing to perform undo operations. When editing is finished, the edited image is automatically saved under the name of the input image extended with the suffix “_ed”. Additionally, the ROIs and information on the application of the ROIs (order of ROIs, add or remove function applied) is stored in a folder with the name of the edited image. If in CiliaQ Preparator the option to create a background-removed image and to duplicate the image channel was selected, CiliaQ Editor can also perform add operations in background-removed images. Here, the information on the intensity of the added voxels is derived from the duplicate image channel that was not segmented.

#### 4.8.2 CiliaQ (version v0.1.4)

CiliaQ contains a “multi-task manager” like CiliaQ Preparator to allow batch-processing. Processing is completely automated. All preferences for the processing are specified by the user when the plugin is launched (a list of preferences for the data sets presented here is shown in Table 2).

One channel is specified that contains the segmentation of the image into cilia and background (referred to as “reconstruction channel”). In this channel, cilia voxels are detected as voxels with an intensity above 0. Initially, connected cilia voxels are merged into objects using a custom 3D implementation of a Flood-Fill-Algorithm. For all Flood-Fill methods, CiliaQ offers two variants of Flood-Filling, i.e. straight filling only or straight and diagonal filling (Fig. 1E). If an object contains less voxels than a user-defined size threshold it is excluded from the analysis and the corresponding voxels are set to 0 in the channel image. All objects passing the size threshold are technically considered individual cilia. For time-lapse images, an additional, custom 4D implementation of a Flood-Fill-algorithm is used to link objects from different time steps. Each resulting object contains the voxels belonging to one individual cilium over time. If selected by the user, results for objects touching the image borders will not be included in the output files.

For additional segmented channels in the image (if used as “intensity A” or “intensity B” channel), CiliaQ also offers to perform the custom, 3D Flood-Fill method to remove objects below a user-defined size threshold in that channel.

Next, voxel-based parameters are determined for each object as described in Table 1.

Next, each object is skeletonized. To this end, for each time step individually, an 8-bit image is created where all voxels contained in the object are set to an intensity of 255 a.u. and all other voxels are set to 0 a.u. Next, the image is 3-fold scaled in all three dimensions using the scale function implemented in ImageJ [53, 54], employing Bilinear interpolation. Next, a Gaussian blur is applied to the image using the “Gaussian Blur” function of ImageJ [53, 54]. The sigma of the blur is defined by the user. The 3-fold of the specified sigma is applied, as the image is 3-fold scaled. Next, the image is skeletonized using a custom implementation of the plugin Skeletonize3D_ [37] and the skeleton is analyzed using a custom implementation of the plugin AnalyzeSkeleton_ [37]. Based on that skeleton analysis, skeleton based parameters are determined as described in Table 1.

Lastly, CiliaQ saves a number of output files, such as the filtered image and a text file containing the preferences and all results. For more information on the output files, see the user guide on the CiliaQ GitHub repository (https://github.com/hansenjn/CiliaQ). CiliaQ offers to generate 3D visualizations of the filtered image, of the detected skeletons, and for each individual cilium. To this end, CiliaQ uses the filtered image, an image with all largest shortest path points, and individual cilium images, respectively. The latter type of image is created as explained for the skeletonization step above. Next, for 3D visualizations, a customized version of the “Volume Viewer” function from FIJI [63] is implemented into CiliaQ.

#### 4.8.4 Hardware requirements

Any computer that can run ImageJ [53, 54] or FIJI [63] is suited to use CiliaQ. The speed of the analysis is mainly determined by the speed of the processor. The memory (RAM) of the computer should be sufficiently large to load an individual image to be analyzed into the memory two-times. Only 3D visualizations, an optional function of CiliaQ, uses the graphics card of the computer.

### 4.9 Software

Data analysis and statistical analysis was performed in Excel (Microsoft Office Professional Plus 2013, Microsoft), GraphPad Prism (Version 8.1.2, GraphPad Software, Inc.), R (Version 3.6.2, The R Foundation for Statistical Computing), RStudio (Version 1.2.5033, RStudio, Inc.), and MATLAB (Version 2018, The MathWorks, Inc.). All image processing and analysis was performed in ImageJ (Version v1.52i, U.S. National Institutes of Health, Bethesda, Maryland, USA). Plots and Figures were generated using GraphPad Prism (Version 8.1.2, GraphPad Software, Inc.), MATLAB (The MathWorks, Inc.), Adobe Illustrator CC (Version 2018, Adobe Systems, Inc.), and Affinity Designer (Version 1.8.4, Serif (Europe), Ltd.). ImageJ plugins were developed in Java, with the aid of Eclipse Mars.2 (Release 4.5.2, IDE for Java Developers, Eclipse Foundation, Inc., Ottawa, Ontario, Canada).

### 4.10 Code and software availability statement

The CiliaQ workflow involves three java-based ImageJ plugins that are freely available online on the CiliaQ GitHub repository (https://github.com/hansenjn/CiliaQ). The GitHub repository also contains a user guide. Like ImageJ, the CiliaQ plugins are open-source but its application does not require coding knowledge, allowing its flexible use by researchers of diverse backgrounds. The GitHub repository also allows to post issues, whereby, for example, ideas for new functions or parameters can be proposed.

## Acknowledgement

The project was supported by grants from the Deutsche Forschungsgemeinschaft (DFG): SPP1726: grant WA3382/3-1 (to DW) and under Germany’s Excellence Strategy – EXC2151 – 390873048 (to DW), by the Boehringer Ingelheim Fonds (to JNH), and by a Samarbeidsorganet Helse Midt-Norge grant (to NJY).

We thank Jens-Henning Krause for technical support and the fish facility support team at the Kavli Institute for Systems Neuroscience.

## Author contribution statement

J.N.H. and D.W. conceived the project. J.N.H., N.J.Y., and D.W. designed experiments. J.N.H., B.S., N.J.Y performed experiments. J.N.H. and S.R. developed software. J.N.H., S.R., and N.J.Y. performed data analysis. J.N.H. drafted the manuscript. All authors revised and edited the manuscript. J.N.H, N.J.Y., D.W. acquired funding.

